# VEGF-C-mediated Cardiac Lymphangiogenesis Promotes Inflammation Resolution in Autoimmune Acute Myocarditis in Mice

**DOI:** 10.1101/2025.05.25.656051

**Authors:** Nanako Nakanishi, Shiro Nakamori, Shigeru Hara, Keiki Nagaharu, Ryo Matsuyama, Soyoka Fujita, Ryuji Okamoto, Kaoru Dohi, Michiaki Hiroe, Kyoko Imanaka-Yoshida, Kazuaki Maruyama

## Abstract

**BACKGROUND:** Acute myocarditis is an immune-mediated inflammatory disease characterized by myocardial inflammation and edema. Cardiac lymphatic vessels are essential for fluid clearance and immune regulation; however, their role in modulating inflammation in autoimmune myocarditis remains largely unclear.

**METHODS:** We investigated the impact of cardiac lymphangiogenesis using a murine model of experimental autoimmune myocarditis induced by cardiac myosin peptide immunization. Human autopsy samples were analyzed for lymphatic expansion. Mice were treated with VEGF-C C156S, a VEGFR3-specific agonist, beginning one week after immunization. Cardiac lymphangiogenesis, myocardial edema, inflammatory infiltration, fibrosis, and cardiac function were evaluated by immunohistochemistry, echocardiography, gene expression analysis, and water content measurements.

**RESULTS:** The VEGF-C treatment accelerated cardiac lymphangiogenesis, enhanced lymphatic drainage, reduced myocardial edema, and attenuated inflammatory cell infiltration and fibrosis. Echocardiography showed the preservation of left ventricular function. VEGF-C selectively decreased the accumulation of iNOS⁺ inflammatory macrophages without broadly suppressing T cells or reparative macrophages. Bulk RNA sequencing confirmed the down-regulation of inflammatory gene signatures associated with macrophage activation.

**CONCLUSIONS:** The early stimulation of cardiac lymphangiogenesis by VEGF-C promotes the resolution of inflammation, reduces myocardial injury, and preserves cardiac function in autoimmune myocarditis. Targeting the cardiac lymphatic system may represent a promising therapeutic strategy for acute myocarditis.

## Introduction

Myocarditis is an immune-mediated inflammatory disease of the myocardium that is commonly triggered by viral infections, medications, or autoimmune responses^1–4^. In recent years, acute myocarditis has gained increased attention due to its association with immune checkpoint inhibitor–related adverse events^5–8^, as well as emerging infectious diseases, such as COVID-19 and the widespread use of mRNA-based vaccines^9–11^. Histologically, it is characterized by inflammatory cell infiltration and myocardial injury, including myocyte degeneration and necrosis^12,13^. In the acute phase, extensive inflammation and increased vascular permeability lead to myocardial edema—a hallmark feature that serves as a key diagnostic criterion in cardiac magnetic resonance imaging^14,15^.

Acute myocarditis, typically presenting within one month of symptom onset, is classified by its etiology, disease phase, severity, clinical presentation, and histopathology. Histopathologically, myocarditis is further categorized by the predominant infiltrating immune cell type—lymphocytic, eosinophilic, giant cell, or granulomatous—reflecting its immunological heterogeneity^1,12^. Although steroid therapy is effective for eosinophilic and giant cell myocarditis, it shows inconsistent benefits for lymphocytic myocarditis, the most common form^12,16^. As a result, current management is largely supportive, focusing on hemodynamic stabilization and bridging to recovery^17^, underscoring the need for early interventions that suppress inflammation, reduce edema, and prevent fibrotic remodeling.

The cardiac lymphatic system plays a pivotal role in maintaining cardiac homeostasis by facilitating the clearance of interstitial fluid, solutes, and metabolic waste back into the circulation^18–20^. Beyond fluid regulation, cardiac lymphatics actively modulate immune responses following cardiac injury^21–24^. They achieve this by recruiting and evacuating infiltrating immune cells from the myocardium and by transporting cytokines, autoantigens, and pathogen-derived molecules to the draining lymph nodes (dLNs), thereby regulating distal immune activation^25^. Moreover, recent studies suggest that lymphatic endothelial cells (LECs) themselves exert immunoregulatory functions, directly contributing to the control of cardiac inflammation^26–28^. Accordingly, increasing attention has been directed towards the cardioprotective functions of lymphatic vessels in cardiovascular diseases.

In this study, we investigated the regulatory role and functional consequences of lymphangiogenesis during cardiac inflammation using a murine model of experimental autoimmune myocarditis (EAM). This model, induced by cardiac myosin peptide combined with complete Freund’s adjuvant (CFA), recapitulates the key features of human lymphocytic myocarditis. In control animals, lymphangiogenesis peaks 3 weeks (3W) after the initial peptide immunization, corresponding to maximal immune cell infiltration. To examine the impact of enhanced lymphangiogenesis, we administered VEGF-C Cys156Ser, a variant that selectively binds to VEGFR3 on the lymphatic endothelium, starting from 1W post-immunization. This early intervention accelerated lymphatic expansion and significantly reduced myocardial edema, inflammatory infiltration, and subsequent fibrosis, ultimately preserving cardiac function. These results highlight cardiac lymphatics as a novel therapeutic target for modulating inflammation and improving the outcomes of acute myocarditis.

## Results

### Lymphangiogenesis is Observed in the Epicardium of Human Hearts with Acute Myocarditis

To characterize the pathological features and lymphatic vessel distribution in human acute myocarditis, we retrospectively analyzed autopsy specimens from four patients who died of the disease at Mie University. These specimens were compared to those from the hearts of individuals who died of non-cardiac causes (**Supplemental Figure 1**). Clinical characteristics are summarized in **Table 1**. Among myocarditis cases, two were diagnosed as giant cell myocarditis and two as lymphocytic myocarditis. None of the patients had a history of immune checkpoint inhibitor therapy, COVID-19 infection, or mRNA-based vaccination. In control hearts, no significant pathological abnormalities were observed in the heart. Podoplanin⁺ lymphatic vessels were identified primarily in the epicardium, with occasional small vessels in the subendocardium, typically adjacent to blood vessels (**Supplemental Figure 1A–C’**). Lymphocytic myocarditis cases showed lymphocytic infiltration accompanied by focal areas of granulation tissue, reflecting ongoing tissue remodeling. Inflammation in the epicardium was associated with the expansion of podoplanin⁺ lymphatic vessels, whereas no marked lymphangiogenesis was observed in endocardial regions (**Supplemental Figure 1D–F’**). A quantitative analysis of podoplanin⁺ lymphatic vessel numbers and the frequency of dilated vessels revealed that both parameters were significantly higher in the epicardium, but not in the endomyocardium, of acute myocarditis cases than in that of controls (**Supplemental Figure 1G and H**).

These results suggest that reactive lymphangiogenesis and vessel dilation occur predominantly in epicardial regions of the human heart during acute myocarditis, in association with inflammatory cell infiltration.

### Lymphatic Expansion Peaks Concurrently with Inflammation in EAM

To assess lymphatic remodeling during acute myocarditis, we induced EAM in 10-W-old male BALB/c mice by immunizing with the α-myosin heavy chain (α-MyHC) peptide emulsified with CFA. Hearts were collected 0, 1, 2, 3, and 4W post-immunization (**Figure 1A**). At 0 and 1W, the cardiac structure remained intact with minimal inflammatory infiltration or fibrosis (**Figure 1B–E′, L-O, Supplemental Figure 2A–B″**). By 2W, focal cardiomyocyte dropout and interstitial infiltration by inflammatory cells became apparent (**Figure 1F, F′**). Most infiltrating cells were CD68⁺ macrophages, accompanied by CD3⁺ T cells—predominantly CD4⁺ helper T cells—with fewer CD8⁺ cytotoxic T cells (**Figure 1F″–F‴‴, L–O**). FoxP3⁺ regulatory T cells and CD20⁺ B cells were also detected in small numbers. CD11c⁺ cells, which include dendritic cells and macrophages, were present at similar levels to CD4⁺ T cells (**Supplemental Figure 2C–C″, F–H**). At this stage, immature collagen fibers positive for Picrosirius red began to appear (**Figure 1G, G′**). At 3W, inflammation peaked and was accompanied by increased immune cell infiltration, progressive cardiomyocyte loss, and enhanced interstitial fibrosis (**Figure 1H–I′, L–P**). By 4W, inflammation subsided and fibrotic changes became more pronounced. Collagen fibers appeared more densely and uniformly stained, indicative of fibrosis maturation (**Figure 1J–K′, L–P; Supplemental Figure 2E–E″, F–H**). These results reveal a clear temporal sequence in EAM, with inflammation and tissue remodeling peaking at 3W and partially resolving by 4W.

**Figure 1.**
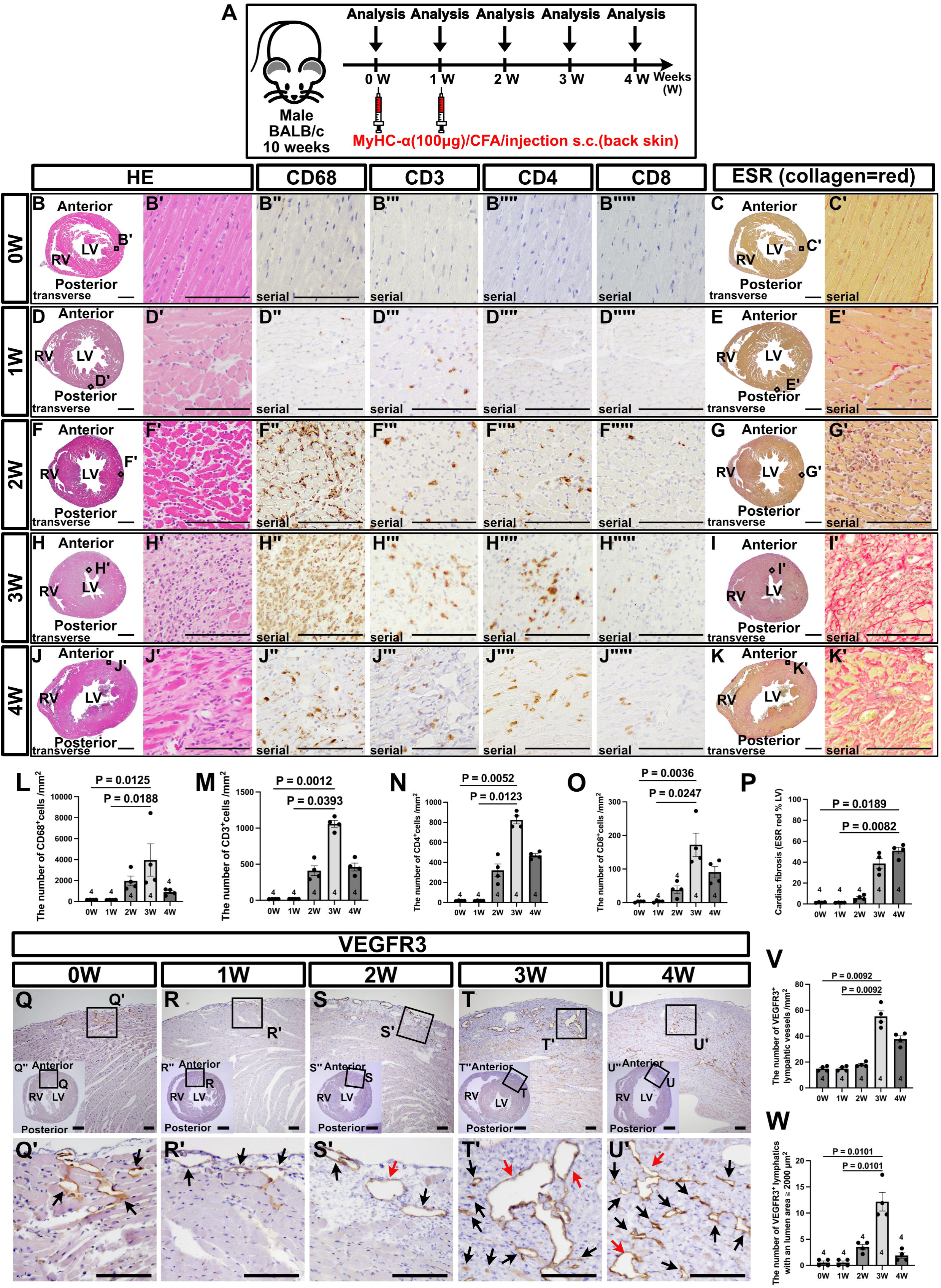
Epicardial lymphangiogenesis peaks at the inflammatory phase in a murine model of autoimmune acute myocarditis. (**A**) Experimental design for the induction and analysis of experimental autoimmune myocarditis. (**B–K**) Transverse histological image analysis of heart sections from the indicated time points stained with hematoxylin and eosin (HE) for general morphology, immunostained for CD68 (macrophages), CD3 (T cells), CD4 (helper T cells), and CD8 (cytotoxic T cells), and Elastica Picrosirius Red (ESR) staining for fibrosis (collagen = red). (**L–O**) Quantitative analysis of inflammatory cells. (**P**) Quantification of cardiac fibrosis (ESR red area as a % of the left ventricular myocardium) at each time point. (**Q–U**’) Representative VEGFR3 immunostaining of heart sections at 0, 1, 2, 3, and 4 weeks, highlighting VEGFR3⁺ lymphatic vessels. Black arrows indicate lymphatic vessels, and red arrows denote those with a larger diameter. (**V, W**) Quantitative analysis of VEGFR3^+^ lymphatic vessels. Each dot represents data from an individual mouse. Scale bars: 1 mm (**B, D, F, H, J, C, E, G, I, K, Q’’, R’’, S’’, T’’, U’’**) and 100 μm (**B’-B””’, D’-D””’, F-F””’, H’-H””’, J’-J””’,C’, E’, G’, I’, K’**). Statistical analyses were performed using the Kruskal–Wallis test followed by Dunn’s multiple comparison test. Error bars represent the mean ± standard error of the mean (SEM).

To further characterize the dynamics of lymphangiogenesis during acute myocarditis, we performed immunostaining for vascular endothelial growth factor receptor 3 (VEGFR3), a marker of LECs. At 0 and 1W, VEGFR3⁺ lymphatic vessels were sparse and mainly located beneath the epicardium, with few being present in endocardial regions (**Figure 1Q–R′, V, W**). As EAM progressed, VEGFR3⁺ lymphatic vessels emerged within inflamed and damaged myocardial areas (**Figure 1S–U′**). By 2W, moderately enlarged lymphatic vessels became apparent (**Figure 1S, S′, V, W**). At 3W, large-diameter and small-caliber vessels both increased in number, consistent with active lymphangiogenesis (**Figure 1T, T′, V, W**). At 4W, as inflammation resolved, the number of VEGFR3⁺ vessels declined, particularly those with larger diameters (**Figure 1U–W**), suggesting the transient and inflammation-driven expansion of the lymphatic network.

To investigate the molecular drivers of this lymphatic response, we assessed the myocardial mRNA expression of key lymphangiogenic factors. *Vegfa* expression progressively decreased after disease onset, whereas *Vegfc* expression peaked at 3W, corresponding to the peak of lymphatic expansion (**Supplemental Figure 2I, J**). These results implicate VEGF-C, rather than VEGF-A, as a major regulator of inflammation-associated lymphangiogenesis during acute myocarditis.

### Sex Differences in the Severity of EAM

Acute myocarditis is reported to occur more frequently in males than in females, based on both clinical and experimental studies^17,29^. To evaluate sex-based differences in our model, we induced EAM in 10-W-old female BALB/c mice using the same immunization protocol (**Supplemental Figure 3A**). A histological analysis revealed that all male mice developed myocardial inflammation and injury, whereas approximately 25% of female mice did not exhibit detectable myocarditis (**Supplemental Figure 3B**). Among female mice that developed inflammation, the extent of immune cell infiltration was consistently smaller than in males across all time points from 2 to 4W, and inter-individual variability was markedly greater (**Supplemental Figure 3C, D**). Although CD68⁺ macrophages and CD11c⁺ cells—representing both dendritic cells and macrophages—were detected, their numbers were generally lower in females, with no significant temporal changes. Similarly, ESR⁺ fibrotic areas showed no consistent time-dependent changes (**Supplemental Figure 3E–R**).

The underlying mechanisms for this reduced and variable disease severity in females remain unclear. However, due to the lower penetrance and higher variability observed in female mice, all subsequent experiments were conducted using males to ensure consistency and reproducibility in disease induction.

### VEGF-C C156S Promotes Cardiac Lymphangiogenesis in EAM

In our EAM model, cardiac lymphatic expansion peaked at 3W post-immunization, coinciding with maximal inflammatory cell infiltration and elevated *Vegfc* mRNA expression. To assess the therapeutic potential of earlier interventions, VEGF-C C156S (hereafter referred to as VEGF-C), a selective VEGFR3 agonist, was administered intraperitoneally at the time of the second peptide injection. Its effects on cardiac lymphangiogenesis were evaluated (**Figure 2A, B**). Whole-mount VEGFR3 immunostaining of the dorsal heart surface revealed increased lymphatic sprouting and slightly enlarged vessel diameters at 2W in VEGF-C–treated mice, while the number of branch points did not significantly differ from phosphate-buffered saline (PBS)-treated controls (**Figure 2C–D′, I–K**). By 3W, no significant differences were observed in sprouting, branching, or diameters between groups (**Figure 2E-F’, I-K**). At 4W, the VEGF-C treatment resulted in a higher number of branch points (**Figure 2G–H′, I–K**). Similar results were observed on the ventral surface (**Supplemental Figure 4A-I**). Despite these morphological changes, the total area occupied by VEGFR3⁺ lymphatic vessels remained similar between groups throughout the 2–4W period (**Supplemental Figure 4J–M**). In transverse sections, the VEGF-C treatment led to increased numbers and larger diameters of VEGFR3⁺ vessels at 2W (**Figure 2L–O**). By 3W, the abundance of large-diameter vessels was reduced in VEGF-C–treated hearts (**Figure 2P-S**), and vessel number and size by 4W were both lower than those at earlier time points (**Figure 2T–W**). High-dose VEGF-C produced similar results (**Supplemental Figure 5A–V**). Ki-67 immunostaining showed an increased percentage of proliferating LECs in VEGF-C–treated hearts at 2W, but no significant differences at 3 or 4W (**Supplemental Figure 4N–T**). In contrast, in mice not subjected to EAM (Peptide−), the VEGF-C treatment did not significantly affect lymphatic sprouting or branching between 2 and 4W. Although a modest increase in vessel diameter was noted at 2W, there were no significant differences in VEGFR3⁺ vessel numbers or Ki-67⁺ LECs in whole-mount or tissue section analyses (**Supplemental Figure 6A-R**).

**Figure 2.**
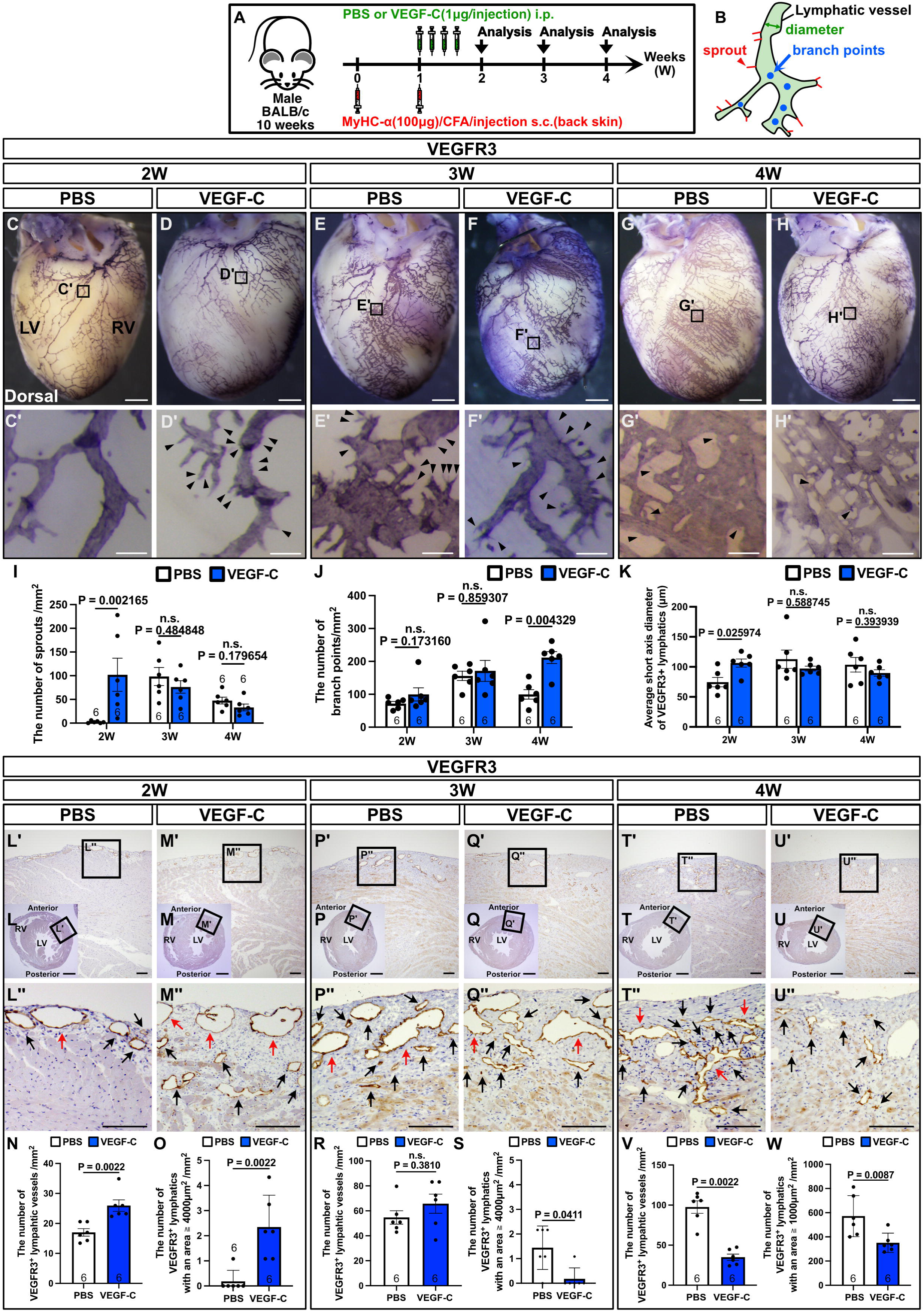
The VEGF-C treatment promotes cardiac lymphangiogenesis in experimental autoimmune myocarditis. (**A**) Experimental timeline for the VEGF-C treatment and subsequent analysis. (**B**) Schematic illustration of parameters assessed in the lymphatic vessel analysis: sprouts, branch points, and vessel diameters. (**C–H′**) Whole-mount VEGFR3 immunostaining of dorsal heart surfaces at 2, 3, and 4 weeks post-immunization in PBS- or VEGF-C–treated mice. Black arrowheads indicate the sprouting of lymphatic vessels. (**I–K**) Quantification of lymphatic morphometric parameters (sprouts, branch points, and diameters) based on whole-mount VEGFR3 staining. (**L–M″, P–Q″, T–U″**) VEGFR3 immunohistochemistry on transverse heart sections from PBS- and VEGF-C–treated mice. Red arrows indicate enlarged lymphatic vessels; black arrows indicate smaller vessels. (**N, O, R, S, V, W**) Quantification of VEGFR3⁺ lymphatic vessels in heart sections. (**N, R, V**) Total number of VEGFR3⁺ lymphatic vessels per mm^2^. (**O, S, W**) Number of VEGFR3⁺ vessels exceeding a predefined luminal area threshold (μm^2^), specific to each time point. Each dot represents an individual mouse. Scale bars: 1 mm (**C, D, E, F, G, H, L, M, P, Q, T, U**) and 100 μm (**C′–H′, L′–L″, M′–M″, P′–P″, Q′–Q″, T′–T″, U′–U″**). Statistical comparisons were performed using the Mann–Whitney U test. Error bars indicate the mean ± standard error of the mean (SEM).

### The VEGF-C Treatment Attenuates Cardiac Edema and Preserves Cardiac Function

We next evaluated the functional effects of the VEGF-C treatment in mice with EAM. An echocardiographic analysis showed that VEGF-C–treated mice maintained better cardiac function than PBS-treated controls. Although the left ventricular internal diameter at diastole (LVIDd) was similar between groups, the left ventricular internal diameter at systole (LVIDs) was significantly preserved in the VEGF-C group at 4W post-immunization (**Figure 3A–C**). In addition, the ejection fraction (EF) and fractional shortening (FS) were both consistently higher in VEGF-C–treated mice from 2 to 4W (**Figure 3D, E**). In the high-dose VEGF-C (VEGF-C HD) group, these improvements were even more pronounced, with significantly better LVIDs, EF, and FS values than that of the controls (**Supplemental Figure 7A–E**). Interventricular septal thickness at diastole and systole (IVSd and IVSs), as well as left ventricular posterior wall thickness at diastole and systole (LVPWd and LVPWs), were also better preserved in VEGF-C–treated mice at 2W, with reduced thinning being observed by 4W (**Figure 3F–I, Supplemental Figure 7F–I**). A gross morphological inspection supported these results, revealing visibly reduced cardiac edema in VEGF-C–treated hearts (**Figure 3J**). Consistently, lymphatic drainage function assessed by Evans blue dye clearance was significantly improved at 2W (**Figure 3K**). Although body weight did not significantly differ between groups, heart weight was significantly lower in VEGF-C–treated mice at 3W (**Figure 3L, M**). Furthermore, myocardial water content was significantly reduced at 2 to 4W, indicating the effective attenuation of cardiac edema **(Figure 3N)**. In contrast, serum cardiac troponin I (cTnI) levels did not significantly differ between PBS- and VEGF-C–treated mice throughout the observation period **(Figure 3O)**.

**Figure 3.**
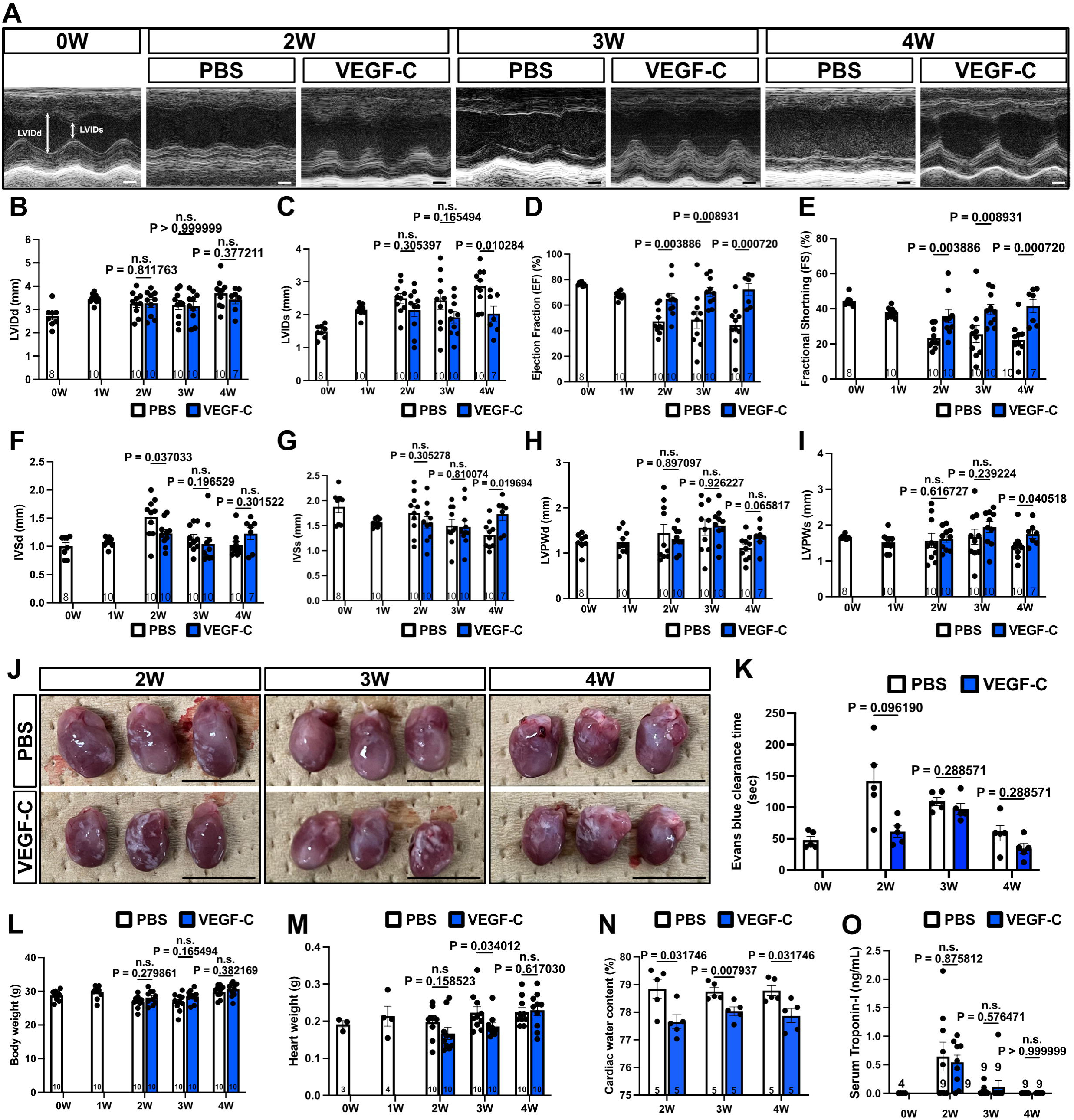
The VEGF-C treatment ameliorates myocardial edema and preserves cardiac function in experimental autoimmune myocarditis. (**A**) Representative M-mode echocardiographic images of the left ventricle (LV) in PBS- and VEGF-C–treated mice at baseline (0W) and at 2, 3, and 4 weeks post-immunization. LVIDd (diastolic LV internal diameter) and LVIDs (systolic LV internal diameter) are indicated. (**B–I**) Quantification of echocardiographic parameters: (**B**) LVIDd, (**C**) LVIDs, (**D**) ejection fraction (EF), (**E**) fractional shortening (FS), (**F**) interventricular septal thickness at diastole (IVSd), (**G**) interventricular septal thickness at systole (IVSs), (**H**) LV posterior wall thickness at diastole (LVPWd), and (**I**) LV posterior wall thickness at systole (LVPWs). (**J**) Representative macroscopic images of hearts harvested at 2, 3, and 4 weeks in PBS- and VEGF-C–treated mice. (**K**) Evans blue dye clearance time, indicating lymphatic drainage efficiency. (**L–N**) Quantitative analysis of body weight (**L**), heart weight (**M**), and cardiac water content (**N**) at each time point. (**O**) Serum cardiac troponin I levels. Each dot represents data from an individual mouse. Scale bars: 1 cm (**A**) and 1 mm (**J**). Statistical analyses were performed using the Mann–Whitney U test. Error bars represent the mean ± standard error of the mean (SEM).

### The VEGF-C Treatment Reduces Infiltrating Macrophages and Attenuates Inflammatory Responses

To examine the effects of VEGF-C on infiltrating inflammatory cells, we analyzed the dynamics of CD68⁺ macrophages—the predominant immune cell population in the myocardium—and CD3⁺ T cells, which are considered key drivers of disease progression (**Figure 1**). At 2 and 4W, no significant differences in CD68⁺ macrophage density were observed between VEGF-C–treated and control groups. However, at 3W, macrophage infiltration was significantly reduced in the VEGF-C group (**Figure 4A–G**). In contrast, CD3⁺ T-cell infiltration remained unchanged across all time points (**Figure 4H–N**). Fibrosis, assessed by Elastica Sirius Red (ESR) staining, was significantly reduced in VEGF-C–treated hearts at both 3 and 4W (**Figure 4O–U**,), with similar changes being observed in the high-dose VEGF-C group (**Supplemental Figure 7J–AD**). A quantitative PCR (qPCR) analysis of whole-heart RNA showed no significant differences in the expression of *Nppb*, *Il1b*, *Il6*, or *Tnfa* at 2W. However, by 3W, the expression levels of these pro-inflammatory genes were significantly lower in VEGF-C–treated hearts than in controls (**Figure 4V–Y**). Bulk RNA sequencing further revealed 1,405 down-regulated and 206 up-regulated genes in VEGF-C–treated hearts relative to PBS-treated controls (**Figure 4Z**). A gene ontology (GO) analysis of down-regulated genes indicated significant enrichment in pathways related to immune regulation, macrophage differentiation, and inflammatory cytokine signaling (**Figure 4AA**). Since macrophages were the only infiltrating cell type to exhibit clear changes upon the VEGF-C treatment, we focused on macrophage-related genes that were both differentially expressed and functionally relevant. Several key inflammatory mediators—including *Clec2d*, *Il6*, *Tnf*, *Il1b*, *Nfkbiz*, *Csf2ra*, *Tlr1*, and *Cd74*—known to be associated with macrophage activation and inflammatory signaling, were markedly down-regulated following the VEGF-C treatment (**Figure 4AB, AC**).

**Figure 4.**
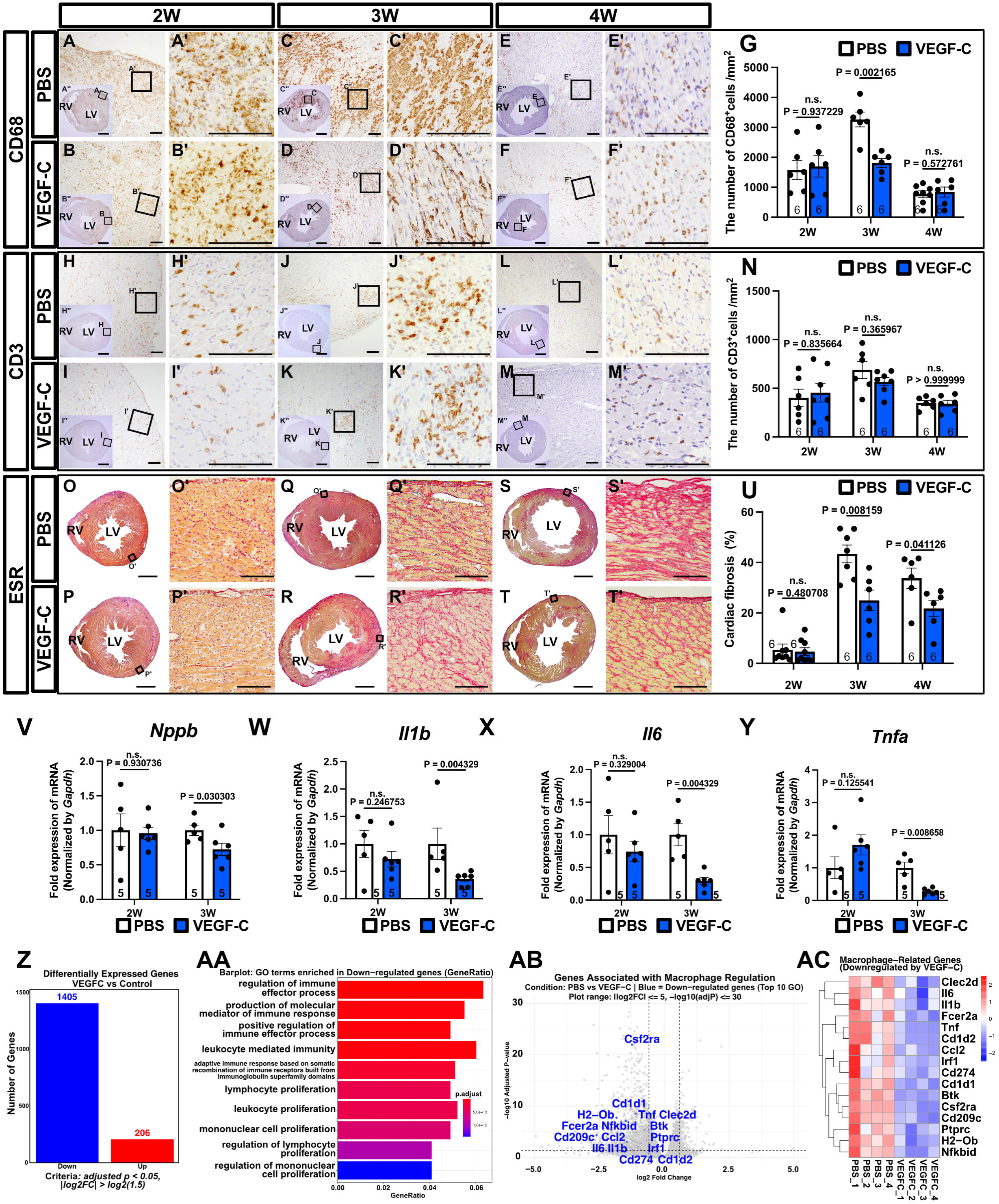
The VEGF-C treatment reduces macrophage infiltration, attenuates myocardial fibrosis, and suppresses inflammatory gene expression in experimental autoimmune myocarditis. (**A–F″, H–M″**) Representative immunohistochemistry for CD68⁺ macrophages (**A–F″**) and CD3⁺ T cells (**H–M″**) in hearts from PBS- and VEGF-C–treated mice at 2, 3, and 4 weeks post-immunization. (**G, N**) Quantification of CD68⁺ macrophages (**G**) and CD3⁺ T cells (**N**) at each time point. (**O–T′**) Representative Elastica Picrosirius Red (ESR) staining showing myocardial fibrosis (collagen = red) in PBS- and VEGF-C–treated mice. (**U**) Quantification of the fibrotic area (ESR⁺ red-stained region as a % of the left ventricular myocardium). (**V–Y**) Quantitative PCR (qPCR) analysis of cardiac mRNA expression for *Nppb* (**V**), *Il1b* (**W**), *Il6* (**X**), and *Tnfa* (**Y**) at 2 and 3 weeks. (**Z**) Summary of differentially expressed genes (DEGs) identified by bulk RNA sequencing between VEGF-C– and PBS-treated hearts (adjusted p <0.05, |log^2^ fold change| >0.584; n = 4 per group). Numbers of up- and down-regulated DEGs are indicated. (**AA**) A gene ontology (GO) enrichment analysis of down-regulated DEGs, highlighting biological processes significantly suppressed by VEGF-C. (**AB**) Enhanced volcano plot showing macrophage-related genes significantly down-regulated in the VEGF-C group. (**AC**) Heatmap visualization of representative macrophage-related genes across individual samples. Each dot represents data from an individual mouse. Scale bars: 1 mm (**A″–F″, H″–M″, O–T**) and 100 μm (**A–F′, H–M′, O′–T′**). Statistical comparisons were performed using the Mann–Whitney U test. Error bars represent the mean ± standard error of the mean (SEM).

### The VEGF-C Treatment Selectively Suppresses Inflammatory iNOS⁺ Macrophages in Autoimmune Myocarditis

To further characterize macrophage heterogeneity during autoimmune myocarditis, we reanalyzed single-cell RNA sequencing (scRNA-seq) data from CD45⁺ leukocytes isolated from the hearts of mice subjected to the same experimental protocol^30^. Consistent with histological results, a large percentage of CD45⁺ cells expressed canonical macrophage markers, including *Cd68*, *Csf1r*, *Cd11b* (*Itgam*), and *Adgre1* (**Figure 5A–C**). An analysis of VEGF receptor expression revealed minimal *Flt4* or *Kdr* expression in CD45⁺ cells, suggesting that VEGF-C acts primarily through LECs rather than directly targeting immune cells (**Supplemental Figure 8A, B**). Furthermore, while a subset of macrophages expressed *Vegfa*, *Vegfc* expression was negligible within CD45^+^ inflammatory cell populations (**Supplemental Figure 8C, D**). Macrophage subclustering based on gene expression profiles identified twelve distinct subsets, including Vegfa⁺ M1-like, Vegfa⁺ M2-like, inflammatory M1-like (*Nos2*⁺, *Tnf*⁺, and *Il1b*⁺), inflammatory monocyte-like (*Ly6c2*⁺ and *Ccr2*⁺), monocyte-derived immature, tissue-repair M2-like (*Arg1*⁺ and *Mrc1*⁺), tissue-resident M2, antigen-presenting (*Cd74*⁺ and *H2-Ab1*⁺), antiviral-responsive (*Isg15*⁺ and *Ifit3*⁺), proliferative (*Mki67*⁺ and *Top2a*⁺), neutrophil-like, and stress-responsive activated clusters (**Figure 5D, E**). The temporal profiling of macrophage subsets across Days 0, 14, 21, and 60 revealed the marked expansion of the inflammatory M1-like and monocyte-derived immature clusters during the inflammatory peak (Days 14–21), followed by a clear contraction during the recovery phase (Day 60) (**Figure 5F**). Feature plots demonstrated that pro-inflammatory markers, such as *Nos2*, *Ccr2*, *Lgals3*, *Arg1*, and *Spp1*, were highly expressed in inflammatory macrophage subsets at peak inflammation, whereas *Mrc1* expression was more diffusely distributed across multiple clusters without strong temporal variations (**Figure 5G**). Since the VEGF-C treatment reduced total macrophage infiltration at 3W (**Figure 4**), we assessed changes in macrophage subsets using immunohistochemistry for key functional markers. A quantitative analysis revealed that VEGF-C selectively reduced the number of iNOS⁺ macrophages—major contributors to myocardial injury—at 3W, whereas the numbers of CCR2⁺, Galectin-3⁺, SPP1⁺, MRC1⁺, and Arginase1⁺ macrophages remained unchanged (**Figure 5H–N**, **Supplemental Figure 8E–O**). These results suggest that VEGF-C preferentially suppresses the accumulation of inflammatory iNOS⁺ macrophages without broadly affecting reparative or regulatory macrophage subsets involved in immune modulation and tissue remodeling.

**Figure 5.**
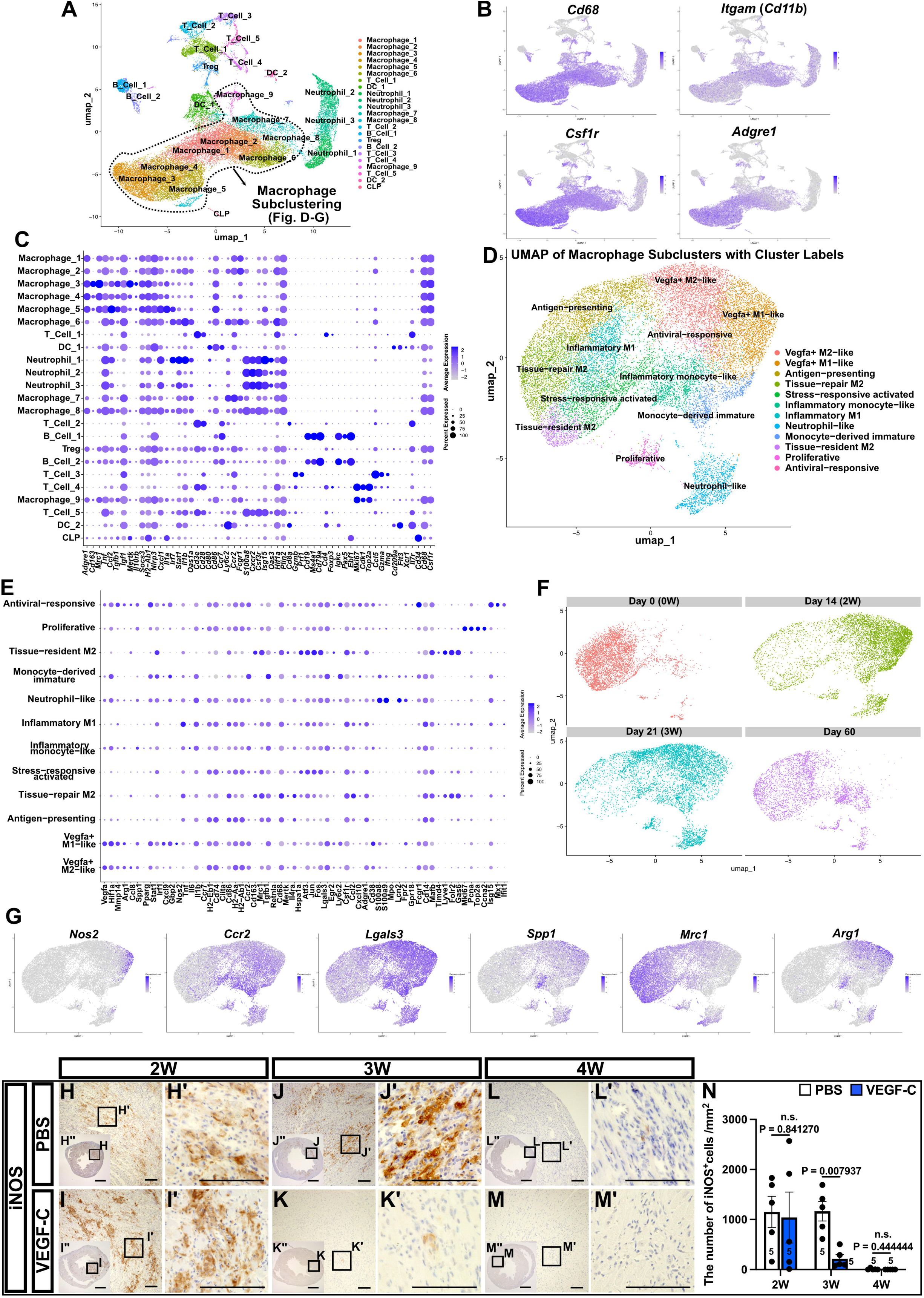
The VEGF-C treatment selectively reduces inflammatory iNOS⁺ macrophages in experimental autoimmune myocarditis. (**A**) Single-cell RNA sequencing (scRNA-seq) analysis of CD45⁺ leukocytes isolated from inflamed hearts during autoimmune myocarditis. UMAP visualization shows overall immune cell clustering. (**B**) UMAP feature plots for the macrophage markers *Cd68*, *Itgam* (Cd11b), *Csf1r*, and *Adgre1*, confirming the predominance of macrophages among infiltrating CD45⁺ cells. (**C**) Dot plot illustrating the expression of key immune-related genes across major immune cell clusters. (**D, E**) UMAP projections of macrophage subclusters, classified into distinct functional phenotypes based on gene expression profiles shown in (**E**). Subsets include Vegfa⁺ M2-like, Vegfa⁺ M1-like, antigen-presenting, tissue-repair M2, stress-responsive activated, inflammatory M1, inflammatory monocyte-like, monocyte-derived immature, neutrophil-like, tissue-resident M2, proliferative, and antiviral-responsive macrophages. (**F**) Temporal dynamics of macrophage subsets from Day 0 (0W) to Day 60, visualized by UMAP. Inflammatory macrophage populations expanded at the peak phase (Days 14–21) and contracted during the recovery phase (Day 60). (**G**) Feature plots showing the expression of representative macrophage functional genes (*Nos2*, *Ccr2*, *Lgals3*, *Spp1*, *Mrc1*, and *Arg1*) over the course of myocarditis. (**H–M″**) Representative immunohistochemistry for inducible nitric oxide synthase (iNOS) in PBS- and VEGF-C–treated hearts at 2, 3, and 4 weeks post-immunization. (**N**) Quantification of iNOS⁺ macrophages in heart sections. The VEGF-C treatment significantly reduced iNOS⁺ macrophages at 3 weeks. Each dot represents data from an individual mouse. Scale bars: 1 mm (**H″–M″)** and 100 μm (**H–M′**). Statistical analyses were performed using the Mann–Whitney U test. Error bars represent the mean ± standard error of the mean (SEM).

## Discussion

In this study, we demonstrated that the stimulation of cardiac lymphangiogenesis by VEGF-C C156S, a selective VEGFR3 agonist, accelerated the resolution of inflammation, reduced myocardial edema and fibrosis, and preserved cardiac function in a murine model of autoimmune myocarditis.

We initially showed that inflammation-associated lymphangiogenesis occurred in autopsy hearts from patients who died of acute myocarditis, where expanded lymphatic networks were observed in inflamed epicardial regions. A similar spatiotemporal pattern was recapitulated in an α-MyHC/CFA-induced EAM model in BALB/c mice, in which inflammation is known to predominantly affect the subepicardial region^31^. In this model, lymphatic expansion peaked at 3W post-immunization, coinciding with maximal immune cell infiltration. This was accompanied by the transient up-regulation of *Vegfc* mRNA and a reduction in *Vegfa* expression, suggesting a temporally regulated, inflammation-driven lymphangiogenic response^32^. Notably, VEGF-C C156S was administered starting on Day 7—approximately 2W prior to the natural peak of *Vegfc* mRNA expression at Day 21—and continued for four consecutive days. This early intervention, rather than prophylactic administration, accelerated lymphangiogenesis and changed the disease course: inflammatory cell infiltration was attenuated, myocardial water content was reduced, and echocardiographic parameters, including EF, FS, and LVIDs, were preserved. These functional improvements were paralleled by reduced myocardial fibrosis, highlighting the therapeutic potential of targeting lymphatic remodeling to mitigate both acute inflammatory injury and subsequent chronic remodeling.

Beyond their structural role in maintaining fluid balance, cardiac lymphatics are increasingly being recognized as active regulators of immune responses during cardiac injury^23,25,33^. Our results suggest that enhancing lymphangiogenesis through VEGF-C is not merely supportive, but actively reshapes the inflammatory microenvironment. Notably, VEGF-C suppressed iNOS⁺ inflammatory macrophages, key drivers of myocardial injury, while preserving CD3⁺ T cell infiltration. A temporal analysis of macrophage subtypes by single-cell RNA sequencing revealed that inflammatory M1-like and monocyte-like clusters predominated during peak myocarditis (Days 14–21) and markedly declined by the recovery phase (Day 60). Immunohistochemistry confirmed that the VEGF-C treatment selectively reduced iNOS⁺ macrophages, primarily associated with the inflammatory M1 subset. In contrast, the densities of CCR2⁺, Galectin-3⁺, SPP1⁺, MRC1⁺, and Arginase1⁺ macrophages—subsets implicated in tissue repair or the amplification of inflammation—were not significantly affected. These results suggest that lymphatic expansion preferentially attenuates pathogenic inflammatory macrophages without broadly suppressing reparative or regulatory subsets.

Previous studies using electron microscopy described cardiac lymphatic vessels as thin-walled structures involved in interstitial fluid clearance, characterized by electron-dense plasma protein deposits uniformly distributed within the lumen^34,35^. Based on these findings, a widespread lymphatic plexus spanning the epicardial to subendocardial layers was initially proposed. However, recent advances in gene regulatory networks governing lymphatic endothelial identity^36–39^ have led to the identification of lymphatic-specific markers such as Prox1, VEGFR3, LYVE1, and podoplanin (D2-40), thereby refining our anatomical understanding of the cardiac lymphatic system^40,41^. In human hearts, lymphatic vessels are now recognized to reside mainly in the subepicardial layer, with sparse extensions along major coronary arteries reaching the endocardium^42,43^. In mice, a similar subepicardial distribution has been observed using markers such as Prox1, LYVE1, and VEGFR3; however, cardiac lymphatic vessels are more specifically aligned along the coronary veins^22^. Despite this venous association, their distribution remains primarily subepicardial, with only minimal extension toward the endocardial surface^44,45^. In acute myocarditis, where inflammation often reaches the subepicardial space, lymphangiogenesis may be induced by inflammatory cytokines (e.g., NF-κB–driven signals) or reflect passive dilation due to increased lymphatic flow^46^. Regardless of the underlying mechanisms, the present results demonstrate that lymphangiogenesis occurs within the subepicardial layer in human acute myocarditis.

Several mechanisms may contribute to the cardioprotective effects of lymphatic expansion. Enhanced lymphatic drainage may reduce local concentrations of pro-inflammatory cytokines, such as IFN-γ and TNF-α, which are essential for sustaining M1 macrophage polarization^24,47,48^. Furthermore, the expansion of the lymphatic network may affect macrophage activation and survival by modulating interstitial pressure, the extracellular matrix composition, and local fluid dynamics^49^. Recent studies using models of acute myocarditis or myocardial infarction highlighted the critical role of macrophage subsets in modulating inflammation and lymphangiogenesis^30,50^. In a transverse aortic constriction model, Heron et al. reported that the blockade of VEGFR3 inhibited lymphangiogenesis without affecting total CD68⁺ macrophage numbers, yet selectively increased iNOS⁺ macrophage infiltration into the heart, exacerbating pathological remodeling^51^. Furthermore, macrophage-derived VEGF-C has been shown to promote lymphangiogenesis and exert cardioprotective effects in both virus-mediated acute myocarditis and infarction models^52,53^.

However, our single-cell RNA-seq reanalysis did not detect *Vegfc* expression in macrophages or other CD45⁺ immune cells. Nevertheless, the bulk mRNA analysis of whole-heart tissue showed that *Vegfc* expression peaked at 3W post-myocarditis induction, suggesting that in our model, VEGF-C was primarily produced by non-immune cells. This interpretation is consistent with previous studies identifying stromal cells^54^, cardiomyocytes^21,55^, and endothelial cells^56^ as potential sources of VEGF-C in the injured heart. Collectively, these results support the notion that macrophages have diverse roles in inflammation and tissue repair, and that the cardioprotective effects of lymphangiogenesis may be mediated, at least in part, by the selective reduction of pro-inflammatory iNOS⁺ (M1-like) macrophages rather than the broad suppression of all macrophage subsets.

Although previous studies showed that VEGFR3 (encoded by Flt4), the receptor for VEGF-C, may be expressed by various cell types under pathological conditions^57,58^, our single-cell RNA-seq reanalysis did not detect *Flt4* expression in macrophages or other CD45⁺ inflammatory cells in this model. These results suggest that LECs are the primary cellular targets of VEGF-C in the context of autoimmune myocarditis. Therefore, the anti-inflammatory effects of VEGF-C may not be mediated by direct effects on immune cells. Instead, LECs appear to be the principal targets, and their functional enhancement may be responsible for the observed therapeutic effects. Nonetheless, while LECs are likely the primary mediators, we cannot entirely exclude the possibility that VEGF-C exerted modest effects on other cardiac cell types, such as cardiomyocytes, vascular endothelial cells, or fibroblasts, although immunostaining in this model did not clearly demonstrate VEGFR3 expression in these cell types.

Beyond immune modulation, the present results highlight the pathophysiological relevance of myocardial edema, which is increasingly being recognized as a direct contributor to cardiac dysfunction. Even a modest 2.5% increase in cardiac water content may reduce cardiac output by 30–40%^60,61^. In this context, lymphatic drainage plays a pivotal role in preserving myocardial performance. In our model, VEGF-C–treated mice exhibited ∼1.5% lower cardiac water content at 2W post-immunization than controls, along with preserved myocardial wall thickness, supporting a functional link between enhanced lymphatic clearance and improved cardiac function.

Sex-based differences in myocarditis were also evident in this model. Male mice exhibited more consistent and severe disease, consistent with previous findings indicating that Th1-skewed immune responses were more pronounced in males and contributed to heightened autoimmune inflammation^29,62^. In contrast, female mice showed greater variability and often failed to develop myocarditis despite identical immunization protocols. This sex-based difference in susceptibility may be partly explained by the immunomodulatory effects of estrogen. Estradiol has been shown to down-regulate pro-inflammatory cytokines, suppress Th1 and Th17 responses, and promote regulatory pathways, thereby conferring protection against autoimmune myocarditis in female mice^29^. Recent studies have reported that male and female mice exhibit innate differences in cardiac lymphatic development, with females having a greater density of lymphatic vessels than males, particularly in the C57BL/6 background^63^. However, in our BALB/c mouse model, we did not observe any marked sex-related differences in cardiac lymphatic vessel distribution. While further investigations into sex-based differences are warranted, understanding why female mice are resistant to developing fulminant acute myocarditis may provide important insights for identifying future therapeutic targets.

In conclusion, the present study provides strong evidence for the early stimulation of cardiac lymphangiogenesis by VEGF-C C156S conferring multifaceted benefits in autoimmune myocarditis, including the attenuation of inflammation, a reduction in myocardial edema, the limitation of fibrosis, and the preservation of cardiac function. The selective reduction of iNOS⁺ macrophages highlights a promising immunomodulatory effect mediated primarily through LEC activation. These results reposition the cardiac lymphatic system from a passive structural component to an active, targetable axis for therapeutic interventions for inflammatory heart disease. Further studies on human tissues and clinical models will be critical to confirm these results and advance the development of lymphatic-targeted therapies towards clinical applications.

## Methods

### Human Samples

Formalin-fixed, paraffin-embedded cardiac tissues were obtained from residual autopsy specimens originally collected for diagnostic purposes. At the time of autopsy, informed consent was obtained from the next of kin, which explicitly included permission for the use of tissues in medical research. This study was approved by the Ethical Review Board of the Graduate School of Medicine, Mie University (approval number: H2024-174). The present study was conducted in accordance with the Ethical Guidelines for Medical and Health Research Involving Human Subjects promulgated by the Japanese government, the principles of the Declaration of Helsinki, and the Department of Health and Human Services Belmont Report.

### Animals and the VEGF-C Treatment Protocol

Male and female BALB/cAJcl mice (10W old) were purchased from CLEA Japan (Tokyo, Japan). All animal experiments were approved by the Mie University Animal Care and Use Committee and were conducted in accordance with institutional guidelines (Approval Nos. 663 and 728). To induce EAM, mice were immunized subcutaneously on Days 0 and 7 with 100 μg α-MyHC (peptide sequence: Ac-RSLKLMATLFSTYASADR; Toray Research Center), emulsified in a 1:1 ratio with PBS and CFA (containing 1 mg/mL *Mycobacterium tuberculosis* H37Ra; Sigma–Aldrich).

Regarding the VEGF-C treatment, mice were randomly assigned to a PBS-treated control group or VEGF-C–treated group. The recombinant human VEGF-C C156S protein (Cat. No. 752-VC/CF; R&D Systems) was administered via intraperitoneal injection at a dose of 1 μg (low dose) or 2 μg (high dose) once daily on Days 7 to 10. The dosing scheme was based on previous studies by Klotz et al^22^. and Lin et al^64^. Since phenotypic changes were evident at the low dose, it was employed as the standard dose in the majority of experiments. Notably, the administration of the high dose (2 μg) enhanced the clarity and consistency of experimental changes.

### Immunohistochemistry and Histology

In section histological analyses, samples were collected and fixed in 2% paraformaldehyde overnight at 4°C, followed by storage in 70% ethanol at 4°C. HE staining and IHC were performed using 3-μm-thick paraffin-embedded sections. Sections for IHC were deparaffinized and rehydrated through a series of xylene and ethanol. In the enzyme-antibody method, endogenous peroxidase activity was blocked using 0.3% hydrogen peroxide (H_2_O_2_) in methanol for 20 min. Antigen retrieval was performed using a pressure chamber with Tris-EDTA buffer (7.4 mM Tris, 1 mM EDTA-2Na, pH 9.0). Sections were then incubated with the following primary antibodies: D2-40 (anti-Podoplanin) (413151, Nichirei Biosciences, at its original concentration (ready to use product)), VEGFR3 (R&D Systems, AF743, 1:150, RRID: AB_355563), CD68 (Cell Signaling Technology, 97778, 1:150, RRID: AB_2928056), CD3 (Abcam, ab5690, 1:150, RRID: AB_305055), CD4 (Abcam, ab183685, 1:150, RRID: AB_2686917), CD8 (Abcam, ab209775, 1:150, RRID: AB_2860566), Foxp3 (Abcam, ab215206, 1:100, RRID: AB_2860568), CD20 (Abcam, ab64088, 1:300, RRID: AB_1139386), CD11c (Cell Signaling Technology, 97585, 1:100, RRID: AB_2800282), iNOS (Cell Signaling Technology, 68186, 1:500, RRID: AB_3662912), Arginase1 (Cell Signaling Technology, 93668, 1:100, RRID: AB_2800207), CCR2 (Abcam, ab273050, 1:150, RRID:A AB_2893307), Galectin-3 (Cell Signaling Technology, 89572, 1:100, RRID: AB_2800111), SPP1 (Cell Signaling Technology, 88742, 1:100, RRID: AB_3107209), MRC1 (Abcam, ab64693, 1:100, RRID:AB_1523912), and Ki67 (Abcam, ab15580, 1:250, RRID: AB_443209). In ESR staining, sections were deparaffinized, rehydrated through graded ethanol, and rinsed in Milli-Q water. Sections were treated with 1% hydrochloric acid in 70% ethanol, stained with Weigert’s resorcin-fuchsin solution for 40–50 minutes to visualize elastic fibers (dark purple-black), and counterstained with Weigert’s iron hematoxylin for 3–5 minutes. After rinsing, sections were stained with Picrosirius Red solution for 15 minutes to highlight collagen fibers in red. In this staining protocol, muscle fibers appear yellow and elastic fibers remain black. Special staining reagents were purchased from Muto Pure Chemicals (Tokyo, Japan). In whole-mount immunostaining, hearts were fixed in 2% paraformaldehyde at 4°C overnight, washed with PBS, and incubated in 0.3% H_2_O_2_ in 100% methanol for 1 hour to block endogenous peroxidase activity. After washing with PBS, tissues were blocked with 2% skim milk and 1% Triton X-100 in PBS. Samples were incubated two to three overnight at 4°C with an anti-VEGFR3 antibody (R&D Systems, AF743, 1:50, RRID: AB_355563). After washing, a HRP-conjugated donkey anti-goat secondary antibody (Abcam, ab6885, 1:500, RRID: AB_955423) was applied, and immunoreactivity was visualized with DAB substrate.

### Echocardiography

Transthoracic echocardiography was performed under 3% isoflurane anesthesia using the Vevo 2100 Imaging System (VisualSonics Inc, Toronto, Canada). The left ventricle (LV) was first visualized in the parasternal long-axis view and then rotated 90 degrees to obtain a short-axis view at the level of the papillary muscles, where M-mode images were acquired to measure LVIDd and LVIDs. LV volumes at diastole (LV Vol;d) and systole (LV Vol;s) were calculated using the following formula: LV Vol = (7.0 / (2.4 + LVID)) × LVID³. EF and FS were calculated from these values as EF (%) = [(LV Vol;d − LV Vol;s) / LV Vol;d] × 100 and FS (%) = [(LVIDd − LVIDs) / LVIDd] × 100, respectively.

### Gravimetry

Cardiac water content was assessed using the wet weight–dry weight method^23^. Hearts were harvested, weighed to obtain the wet weight, and then desiccated for 5 days at 65°C to obtain the dry weight. Cardiac water content (%) was calculated using the following formula: (wet weight − dry weight) / wet weight × 100.

### qPCR

Cardiac tissues were homogenized, pooled, and subdivided for RNA extraction. Total RNA was isolated using Isogen reagent (Nippon Gene, Tokyo, Japan) according to the manufacturer’s instructions, with Gene Packman (Nacalai Tesque, Kyoto, Japan) added to enhance RNA yield. First-strand cDNA synthesis was performed using the Transcription First Strand cDNA Synthesis Kit (Roche Diagnostics, Mannheim, Germany). SYBR Green-based qPCR was conducted using the LightCycler 96 System (Roche, Germany) in combination with the QuantiTect SYBR Green PCR Kit and QuantiTect Primer Assays (Qiagen, Hilden, Germany), following the manufacturer’s protocol. Data were analyzed using LightCycler 96 software (Roche), and gene expression was normalized to the housekeeping gene Gapdh using the 2^−ΔCt^ method. In selected experiments, TaqMan-based qPCR was performed. Total RNA was directly used for amplification with the TaqMan RNA-to-Ct 1-Step Kit (Thermo Fisher Scientific, Waltham, MA, USA) or cDNA was synthesized using the High-Capacity cDNA Reverse Transcription Kit (Thermo Fisher Scientific). Amplification was performed using the QuantStudio 7 Flex Real-Time PCR System (Thermo Fisher Scientific) and TaqMan Gene Expression Master Mix. The following TaqMan probes (Qiagen) were used: *Nppb* (QT00107541), *Vegfc* (QT00104027), *Vegfa* (QT00160769), *Il1b* (QT01048355), *Il6* (QT00098875), *Tnf* (QT00104006), and *Gapdh* (QT01658692). Expression levels were normalized to *Gapdh* and are presented as fold changes relative to the control group, as shown in the corresponding figures.

### Bulk RNA Sequencing

Total RNA was extracted from LECs using Isogen reagent (Nippon Gene, Tokyo, Japan) according to the manufacturer’s protocol. RNA quantity and purity were assessed using a NanoDrop spectrophotometer, and RNA integrity was confirmed using the 2100 Bioanalyzer (Agilent Technologies). Samples with an RNA integrity number >8.0 were subjected to a downstream analysis. Sequencing libraries were prepared at the Research Institute for Microbial Diseases, Osaka University, and sequenced using the Illumina NovaSeq 6000 platform. Raw FASTQ files were processed with Trim Galore (v0.6.10) for adapter trimming and STAR (v2.7.10a) for read alignment. Gene-level read counts were obtained using featureCounts (Subread v2.0.1)^65^. Differential gene expression analysis was performed using DESeq2 (v1.38.3)^66^, and further visualized and explored using the iDEP web application (v2.01)^65^. Genes with an adjusted p-value < 0.05 and an absolute log₂ fold change > 0.584 (corresponding to a 1.5-fold change) were considered significantly differentially expressed. The numbers of upregulated and downregulated genes are summarized in Supplementary Table 1. Gene ontology (GO) enrichment analysis of downregulated genes was conducted using the clusterProfiler R package and visualized as bar plots. To specifically investigate the immunomodulatory effects of VEGF-C, macrophage-related DEGs were extracted by cross-referencing genes annotated in the top 10 downregulated immune-related GO terms with curated macrophage gene sets. These genes were visualized using a custom volcano plot with fixed axis limits (|log₂FC| ≤ 5, –log₁₀ p ≤ 30) to enhance interpretability. Additionally, a heatmap of representative macrophage-regulatory genes was generated using the pheatmap package.

### scRNA-seq

Previously published scRNA-seq data (GEO: GSE142564)^30^ profiling CD45⁺ immune cells isolated from murine hearts at various phases of EAM (Days 0, 14, 21, and 60) were reanalyzed. Raw sequencing data were processed using Cell Ranger (v6.1.2, 10x Genomics) and aligned to the mm10 reference genome. The resulting feature-barcode matrices were imported into R (v4.3.1) and analyzed with the Seurat package (v5.1.0). After quality control filtering to exclude cells with fewer than 1,000 detected genes or >5% mitochondrial gene content, data were normalized using the NormalizeData function (log-normalization), and highly variable genes were identified with FindVariableFeatures. A principal component analysis was performed on scaled data, and the first 10 principal components were used for dimensionality reduction via uniform manifold approximation and projection (UMAP), as well as for shared nearest neighbor graph construction and clustering via the Louvain algorithm (FindClusters, resolution = 0.5). no batch correction or data integration (IntegrateData) was performed across timepoints. All downstream analyses were conducted using the default RNA assay to preserve biological variation between disease stages. UMAP visualizations and differential expression analyses therefore reflect gene expression within the native RNA assay.

Macrophage clusters were identified based on canonical marker expression (*Cd68*, *Itgam*, *Csf1r*, and *Adgre1*), a subset using the subset function, and reanalyzed through the same pipeline to resolve macrophage subclusters. Gene expression was visualized using FeaturePlot and DotPlot, and DEGs within each cluster were identified using FindAllMarkers with the Wilcoxon rank-sum test and Bonferroni correction. Clusters were annotated based on curated marker gene sets: inflammatory M1-like (*Nos2*, *Tnf*, *Il1b*, and *Cxcl9*), antigen-presenting (*Cd74*, *H2-Ab1*, *Cd86*, and *Ciita*), tissue-repair M2-like (*Arg1*, *Mrc1*, *Tgfb1*, and *Cd163*), stress-responsive (*Atf3*, *Jun*, *Fos*, and *Hspa1a*), monocyte-derived immature (*Ly6c2*, *Ccr2*, and *Csf1r*), neutrophil-like (*S100a8*, *S100a9*, and *Mpo*), proliferative (*Mki 67*, *Top2a*, and *Pcna*), and tissue-resident M2-like macrophages (*Folr2*, *Cd209g*, and *Gas6*). In addition, clusters co-expressing Vegfa, a growth factor that, like Vegfc, potentially promotes lymphangiogenesis^67^, along with either M1- or M2-associated genes, were designated as Vegfa⁺ M1-like or Vegfa⁺ M2-like subsets.The temporal dynamics of these subclusters were assessed across disease phases (Days 0–60), and representative marker genes were visualized by UMAP and feature plots to illustrate functional transitions in the macrophage compartment. All analyses were performed using base R and Seurat functions, and graphical outputs were exported in the high-resolution PDF format for figure integration.

### Quantification of Inflammatory Cell Infiltration and the Fibrotic Area

All cardiac specimens were transversely sectioned into three equal parts to enhance the detection sensitivity of myocarditis. From each of the three levels, 3-μm transverse sections were prepared. At a minimum, one hematoxylin and eosin (HE)-stained section and nine serial sections for immunohistochemistry were generated per level. The quantification of inflammatory cells was performed on immunohistochemically stained cardiac sections using a 40× objective lens (total magnification ∼400×). For each mouse, three transverse heart sections were prepared, and two representative sections were selected based on overall histological quality. Within each selected section, two areas with the most intense inflammatory cell infiltration were identified at the mid-ventricular level. In these areas, positively stained cells were manually counted by identifying hematoxylin-stained nuclei within the antibody-stained cells. The average cell count was calculated for the two regions per section, and the final value per mouse was determined by averaging results from both sections.

The fibrotic area was quantified using ESR staining. Images were captured at 2× objective lens (total magnification ∼20×), and among the three sections per animal, that at the papillary muscle level exhibiting the largest red-stained area was selected for analysis. Images were processed in ImageJ (NIH) by converting to RGB stacks. Red-stained collagenous areas were isolated using manual threshold adjustments. The LV myocardium was defined as the region of interest (ROI) using the polygon selection tool, and the percentage of the red-stained area within the LV myocardium was calculated.

The incidence of myocarditis was defined as the percentage of mice showing histological evidence of inflammatory cell infiltration, with or without associated myocardial injury—such as cardiomyocyte necrosis or replacement fibrosis—on HE-stained sections.

### Quantification of Lymphatic Vessels

Whole-mount immunostaining for VEGFR3 was performed on hearts imaged from both the dorsal and ventral surfaces. In VEGFR3-stained whole-mount hearts, lymphatic sprouting was defined as the emergence of thin, fine-caliber protrusions extending from existing lymphatic vessels, whereas branch points were defined as bifurcations involving thicker lymphatic structures, excluding fine sprouts. For each heart, two 500 × 500 μm regions of interest (ROIs) displaying prominent lymphatic sprouting were selected. The number of sprouts and branch points within each ROI was manually counted, normalized per mm², and the average of the two values was used for analysis. To assess vessel calibers, the three largest valved lymphatic segments were selected per sample, and the maximum short-axis diameter between valves was measured in each segment. The average of these three measurements was calculated and used as the representative vessel diameter. The overall lymphatic area (% VEGFR3⁺ area) was calculated by excluding great vessels and applying contrast-limited adaptive histogram equalization (CLAHE) and threshold-based binarization using ImageJ.

In tissue sections stained for VEGFR3, lymphatic vessel density was quantified by counting VEGFR3⁺ vessels in two representative 20× fields (total magnification ∼200×) with the highest lymphatic vessel density within inflammatory lesions from each section. The average of these two fields was first calculated per section, and then the average of two separate tissue sections per animal was used to determine the final value, expressed as vessels per mm². To assess lymphatic vessel dilation, the cross-sectional lumen areas of VEGFR3⁺ vessels were measured using ImageJ in two representative 20× fields (total magnification ∼200×) per tissue section. The mean values from two sections per animal were calculated.

For proliferation analysis, paraffin sections previously used for whole-mount VEGFR3 staining were subsequently immunostained for Ki67. VEGFR3⁺ lymphatic vessels were visualized using a DAB substrate supplemented with nickel chloride (NiCl), resulting in a dark purple-black signal, while Ki67 was developed with standard DAB, yielding a brown nuclear stain. This chromogenic contrast allowed clear identification of Ki67⁺ nuclei located within VEGFR3⁺ vessel walls, while excluding inflammatory cells and lymphatic valve structures. In each sample, 100 LEC nuclei were analyzed, and the percentage of Ki67⁺ LECs was calculated.

### Evans Blue Clearance Assay

To evaluate cardiac lymphatic drainage function, Evans blue dye clearance was assessed in anesthetized mice. Mice were anesthetized with 3% isoflurane, followed by orotracheal intubation and mechanical ventilation to maintain respiration during thoracotomy. After surgically opening the chest wall to expose the beating heart, 20 μL of 0.5% Evans blue (FUJIFILM Wako, 056-04061) solution (in PBS) was injected into the subepicardial surface of the apex using a 29G insulin syringe (TERUMO, SS-10M2913). The time required for the dye to disappear from the subepicardial surface was measured as previously described^68^, and this clearance time was recorded as an index of functional lymphatic transport capacity.

### Serum cTnI Measurement

Serum cTnI levels were measured using an ultra-sensitive mouse cardiac troponin-I ELISA kit (Life Diagnostics, Inc., CTNI-1-US) according to the manufacturer’s instructions. Briefly, blood samples were collected from the inferior vena cava at the time of sacrifice, transferred into microcentrifuge tubes, and allowed to clot at room temperature for 1 hour. Following centrifugation at 1,500 × *g* for 15 minutes, serum was collected and stored at −80°C until analyzed. A standard curve ranging from 0 to 2.5 ng/mL was prepared by serial dilutions of the stock standard (100.1 ng/mL) in assay diluent. All samples and standards (200 μL/well) were loaded onto the ELISA plate and incubated at 25°C for 2 hours. After five washes with 1× wash solution, 100 μL of the HRP conjugate and 100 μL of the diluent were added sequentially and incubated at 25°C for 1 hour with gentle shaking. Plates were washed again and developed with 100 μL of TNB substrate for 20 minutes, followed by the addition of 100 μL of stop solution. Absorbance was measured at 450 nm within 5 minutes using a microplate reader (PerkinElmer, MA, USA). cTnI concentrations were calculated based on the standard curve.

### Statistics and reproducibility

All statistical analyses and data visualization were performed using GraphPad Prism version 10 (GraphPad Software). Data are presented as mean ± standard error of the mean (SEM). Comparisons between two groups were performed using the Mann–Whitney U test. The Kruskal–Wallis test followed by Dunn’s multiple comparison test was used for comparisons involving three or more groups. P values less than 0.05 were considered to be significant. No statistical methods were used to predetermine the sample size. Experiments were not randomized, and investigators were not blinded during data collection, analyses, or outcome assessments.

### Data availability

Bulk RNA sequencing data generated in this study have been deposited in the NCBI Gene Expression Omnibus (GEO) under accession number GEO (pending)

## AUTHORS’ CONTRIBUTIONS

K.M. conceived the study and designed the experiments. N.N. and S.N. performed the majority of the experimental work. The single-cell RNA sequencing reanalysis was conducted by K.N. and K.M. The bulk RNA sequencing analysis was performed by S.H., R.M., and K.M. The manuscript was supervised and critically reviewed by S.N., M.H., K.D., R.O., and K.I.-Y. K.M. coordinated the experimental work, analyzed the data, and wrote the manuscript with input from all authors.

## ACKNOWLEDGMENTS

We extend our sincere gratitude to all laboratory members for their insightful discussions and continued encouragement throughout this study. This work was supported in part by Grants-in-Aid for Scientific Research from the Ministry of Education, Culture, Sports, Science and Technology of Japan (23K15949 and 25K12930 to K.M., and 24K02250 to K.I.-Y.); by the VBIC Research Grant from the Japan Foundation for Applied Enzymology (to K.M.); by research grants from the Takeda Science Foundation, the Ichiro Kanehara Foundation for the Promotion of Medical Sciences and Medical Care, the Kurata Research Grant from the HITACHI Global Foundation, the Research Promotion and Graduate School Reform-Related Grant Project from Mie University, the TERUMO Life Science Foundation, the Sumitomo Foundation, the Astellas Foundation for Research on Metabolic Disorders, and the Kenzo Suzuki Memorial Medical Science Foundation (all to K.M.); and we gratefully acknowledge the NGS Core Facility at the Research Institute for Microbial Diseases, Osaka University, for their support with sequencing.

---

**Supplemental Figure 1. Epicardial lymphatic expansion is observed in human acute myocarditis cases.**

(**A–C’**) Representative hematoxylin and eosin (HE) staining and podoplanin immunohistochemistry of control human hearts without cardiac disease. Black arrows indicate podoplanin⁺ lymphatic vessels in the epicardium (**B′**) and endocardium (**C′**). (**D–F’**) Representative HE and podoplanin staining of hearts from patients with lymphocytic myocarditis. Black arrows indicate podoplanin⁺ lymphatic vessels; red arrows highlight vessels with a larger diameter (**E′, F′**). (**G, H**) Quantification of podoplanin⁺ lymphatic vessels and the number of dilated podoplanin⁺ vessels per 2000 μm^2^. Each dot represents an individual case. Scale bars: 1 mm (**B, E**) and 100 μm (**A, B’, C, C’, D, E’, F, F’**). Statistical analyses were performed using the Mann–Whitney U test. Error bars represent the mean ± standard error of the mean (SEM).

**Supplemental Figure 2. Temporal changes in regulatory T cells, B cells, dendritic cells, and lymphangiogenic gene expression during experimental autoimmune myocarditis.**

(**A–E′′**) Representative immunohistochemistry for Foxp3⁺ regulatory T cells (**A–E**), CD20⁺ B cells (**A′–E′**), and CD11c⁺ cells (**A′′–E′′**) in heart sections collected at 0, 1, 2, 3, and 4 weeks post-immunization. CD11c⁺ cells primarily represent dendritic cells, but may also include activated macrophages. (**F–H**) Quantification of Foxp3⁺, CD20⁺, and CD11c⁺ cells over time.

(**I, J**) Quantitative PCR (qPCR) analysis of *Vegfa* (**I**) and *Vegfc* (**J**) mRNA levels in whole heart tissue at the indicated time points, normalized to *Gapdh*. Each dot represents an individual mouse. Scale bars: 100 μm (**A–E′′**). Statistical analyses were performed using the Kruskal–Wallis test followed by Dunn’s multiple comparison test. Error bars represent the mean ± standard error of the mean (SEM).

**Supplemental Figure 3. Sex-based differences in myocarditis severity in experimental autoimmune myocarditis.**

(**A**) Experimental timeline for myocarditis induction and analysis in female BALB/c mice. (**B**) Incidence of myocarditis (%) at 2, 3, and 4 weeks post-immunization in female and male mice. (**C**) Representative HE-stained heart sections at 3 weeks, illustrating differences in myocardial inflammation (dotted lines) between sexes. (**D**) Quantification of the inflammation area (% of the total myocardial area, combining the left and right ventricles) in female and male mice. (**E–J′**) Representative histological and immunohistochemical staining in female hearts at 2, 3, and 4 weeks post-immunization, including HE, CD68, CD3, CD4, CD8, Foxp3, CD20, CD11c, and Elastica Sirius Red (ESR). (**K–Q**) Quantification of immune cell infiltration in female mice across the indicated time points. (**R**) Quantification of the fibrotic area (ESR⁺ red-stained region) in the left ventricular (LV) myocardium, showing no significant differences among time points. Each dot represents an individual mouse. Scale bars: 1 mm (**C, E, G, I, F, H, J**) and 100 μm (**E′–E⁗, G′–G⁗, I′–I⁗, F′, H′, J′**). Statistical analyses were performed using the Kruskal–Wallis test followed by Dunn’s multiple comparison test. Error bars represent the mean ± standard error of the mean (SEM).

**Supplemental Figure 4. VEGF-C promotes the structural adaptation of cardiac lymphatics during autoimmune myocarditis.**

(**A–F′**) Whole-mount VEGFR3 immunostaining of the ventral heart surface at 2, 3, and 4 weeks post-immunization in PBS- or VEGF-C–treated mice. (**G–I**) Quantification of lymphatic morphological features (sprouts, branch points, and diameters) from whole-mount images. (**J, K**) Representative binarized images of VEGFR3⁺ lymphatic vessels at 3 weeks on the ventral (**J**) and dorsal (**K**) heart surfaces. Red outlines indicate the myocardial area used for quantification. (**L, M**) Quantification of the VEGFR3⁺ lymphatic vessel area (%) on ventral (**L**) and dorsal (**M**) surfaces. (**N–S″**) Paraffin sections prepared after whole-mount staining were subjected to Ki67 immunostaining. VEGFR3⁺ lymphatic structures were developed using a DAB substrate with nickel enhancement, resulting in a purple signal, whereas Ki67 staining appeared brown. (**N–S**) Low-magnification views; (**N′–S′**) enlarged views of boxed regions showing Ki67⁺ lymphatic endothelial cells (black arrows). (**T**) Quantification of proliferating Ki67⁺ LECs, expressed as the percentage of Ki67⁺ nuclei among total LECs. Each dot represents data from an individual mouse. Scale bars: 1 mm (**A–F, N″–S″**) and 100 μm (**A′–F′, N-S, N′–S′**). Statistical analyses were performed using the Mann–Whitney U test. Error bars indicate the mean ± standard error of the mean (SEM).

**Supplemental Figure 5. High-dose VEGF-C administration enhances lymphatic sprouting and dilation in experimental autoimmune myocarditis.**

**(A)** Experimental design. Male BALB/c mice (10 weeks old) were immunized with the α-MyHC peptide and CFA on days 0 and 7 to induce myocarditis. Mice were then randomized to receive high-dose VEGF-C C156S (2 μg/injection, intraperitoneally) or PBS daily from Days 7 to 10. Hearts were collected at 2, 3, and 4 weeks post-immunization. (**B–G′**) Representative whole-mount VEGFR3 immunostaining of dorsal heart surfaces at 2, 3, and 4 weeks; black arrowheads indicate lymphatic sprouts. (**H–J**) Quantification of lymphatic sprouting, branching, and vessel diameter from whole-mount images. (**K–L″, O–P″, S–T″**) Representative VEGFR3 immunohistochemistry of paraffin heart sections. (**K–T**) Low-magnification views; (**K′–T′**) higher-magnification views showing VEGFR3⁺ lymphatic vessels (red arrows indicate enlarged lymphatic vessels; black arrows indicate smaller lymphatic vessels). (**M, N, Q, R, U, V**) Quantification of VEGFR3⁺ lymphatic vessels in tissue sections. Each dot represents an individual mouse. Scale bars: 1 mm (**B–G, K″–T″**) and 100 μm (**B′–G′, K–T, K′–T′**). Statistical analyses were performed using the Mann–Whitney U test. Error bars represent the mean ± standard error of the mean (SEM).

**Supplemental Figure 6. VEGF-C administration does not significantly affect the cardiac lymphatic morphology in non-inflamed hearts.**

**(A)** Experimental design. Male BALB/c mice (10 weeks old) were administered either PBS or VEGF-C C156S (1 μg/injection, intraperitoneally) without myocarditis induction (Peptide(−)). Hearts were collected and analyzed at 2, 3, and 4 weeks post-injection. (**B–G′**) Representative whole-mount VEGFR3 immunostaining of dorsal heart surfaces at 2, 3, and 4 weeks in PBS- or VEGF-C–treated mice. Red lines indicate the short-axis diameters of lymphatic vessels. (**H–M″**) Representative VEGFR3 immunohistochemistry on paraffin heart sections. Red arrows indicate VEGFR3⁺ lymphatic vessels, the green arrow indicates larger VEGFR3^+^ lymphatic vessels. Each dot in the quantification graphs represents an individual mouse. Scale bars: 1 mm (**B–G, H–M**) and 100 μm (**B′–G′, H′–M′, H″–M″**). Statistical analyses were performed using the Mann–Whitney U test. Error bars represent the mean ± standard error of the mean (SEM).

**Supplemental Figure 7. High-dose VEGF-C administration reduces cardiac inflammation and fibrosis in experimental autoimmune myocarditis.**

**(A)** Representative M-mode echocardiographic images of the left ventricle (LV) in PBS- and high-dose VEGF-C (HD)-treated mice at baseline (0W) and at 2, 3, and 4 weeks post-immunization. (**B–I**) Quantification of echocardiographic parameters: (**B**) LVIDd (diastolic LV internal diameter), (**C**) LVIDs (systolic LV internal diameter), (**D**) ejection fraction (EF), (**E**) fractional shortening (FS), (**F**) interventricular septal thickness at diastole (IVSd), (**G**) at systole (IVSs), (**H**) LV posterior wall thickness at diastole (LVPWd), and (**I**) at systole (LVPWs). (**J–O″**) Representative immunohistochemistry for CD68⁺ macrophages at 2, 3, and 4 weeks post-immunization. (**P**) Quantification of CD68⁺ macrophage density. (**Q–V″**) Representative immunohistochemistry for CD3⁺ T cells at 2, 3, and 4 weeks. (**W**) Quantification of CD3⁺ T cell density. (**X–AC′**) Representative Elastica Picrosirius Red (ESR) staining for cardiac fibrosis (collagen = red) at each time point. (**AD**) Quantification of the fibrotic area (ESR⁺ region as a % of the left ventricular myocardium). Each dot represents data from an individual mouse. Scale bars: 1 mm (**A, J″–O″, Q″–V″, X–AC**) and 100 μm (**J–O, J′–O′, Q–V, Q′–V′, X′–AC′**). Statistical analyses were performed using the Mann–Whitney U test. Error bars represent the mean ± standard error of the mean (SEM).

**Supplemental Figure 8. Expression of CCR2, Galectin-3, SPP1, MRC1, and Arginase1 is unchanged by the VEGF-C treatment in autoimmune myocarditis.**

(**A–D**) UMAP feature plots showing the single-cell RNA-seq expression of *Kdr (Vegfr2)*, *Flt4 (Vegfr3)*, *Vegfa*, and *Vegfc* in cardiac CD45⁺ leukocytes isolated from mice with experimental autoimmune myocarditis (EAM). (**E–N′**) Representative immunohistochemistry for CCR2 (**E–F′**), Galectin-3 (**G–H′**), SPP1 **(I–J′**), MRC1 (**K–L′**), and Arginase1 (**M–N′**) in hearts from PBS- and VEGF-C–treated mice at 3 weeks post-immunization. (**O**) Quantification of the number of positive cells for each marker. No significant differences were observed between PBS- and VEGF-C–treated groups. Each dot represents data from an individual mouse. Scale bars: 1 mm (**E′–N′**) and 100 μm (**E–N**). Statistical analyses were performed using the Mann–Whitney U test. Error bars represent the mean ± standard error of the mean (SEM).

