## Supplemental Figure and legends for "VEGF-C-mediated Cardiac Lymphangiogenesis Promotes Inflammation Resolution in Autoimmune Acute Myocarditis in Mice"

Supplemental Figures. 1-8

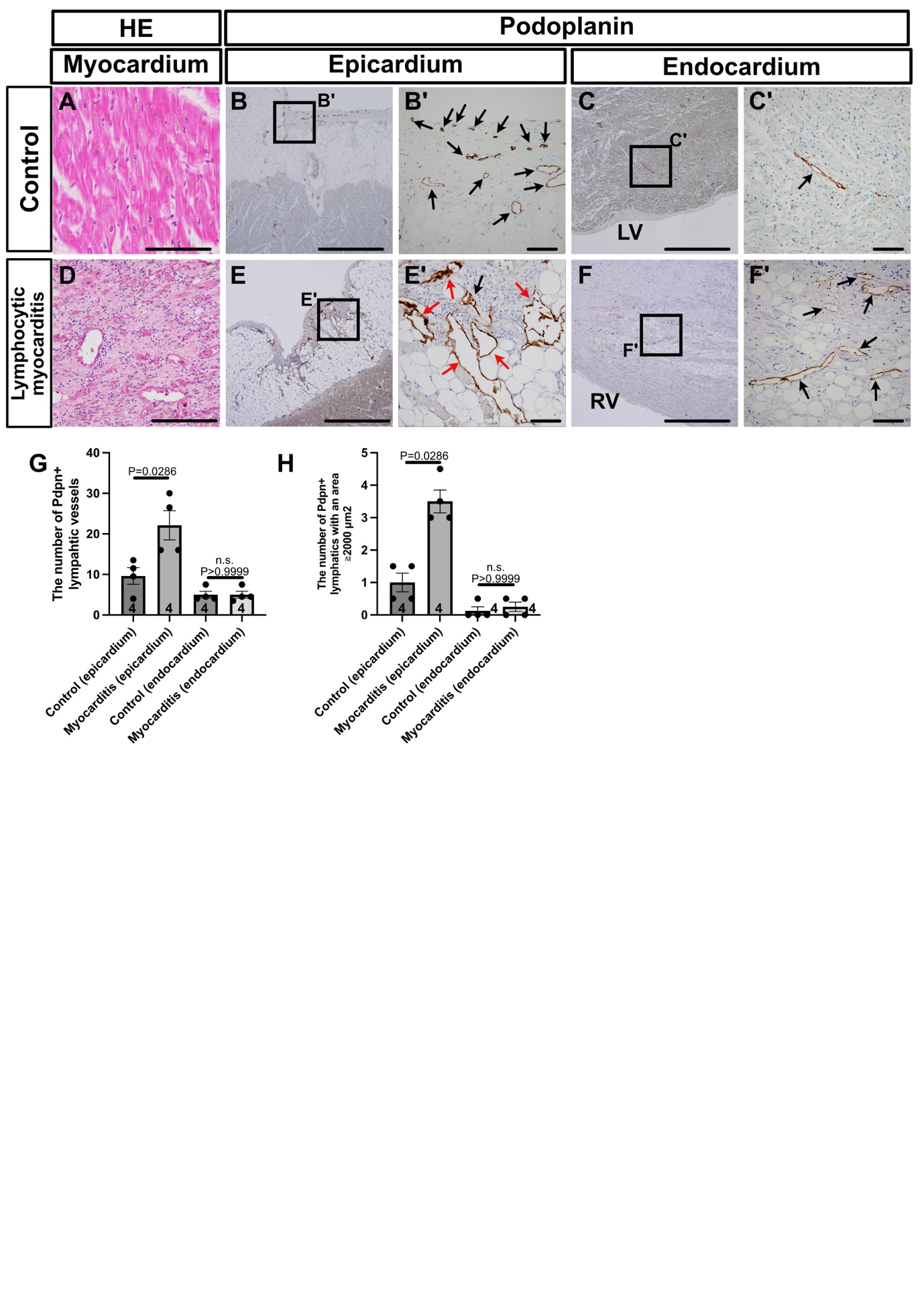
Supplemental Figure 1.

**Supplemental Figure 1. Epicardial lymphatic expansion is observed in human acute myocarditis cases.**

(**A–C'**) Representative hematoxylin and eosin (HE) staining and podoplanin immunohistochemistry of control human hearts without cardiac disease. Black arrows indicate podoplanin⁺ lymphatic vessels in the epicardium (**B′**) and endocardium (**C′**). (**D–F'**) Representative HE and podoplanin staining of hearts from patients with lymphocytic myocarditis. Black arrows indicate podoplanin⁺ lymphatic vessels; red arrows highlight vessels with a larger diameter (**E′, F′**). (**G, H**) Quantification of podoplanin⁺ lymphatic vessels and the number of dilated podoplanin⁺ vessels per 2000 μm^2^. Each dot represents an individual case. Scale bars: 1 mm (**B, E**) and 100 μm (**A, B’, C, C’, D, E’, F, F’**). Statistical analyses were performed using the Mann–Whitney U test. Error bars represent the mean ± standard error of the mean (SEM).

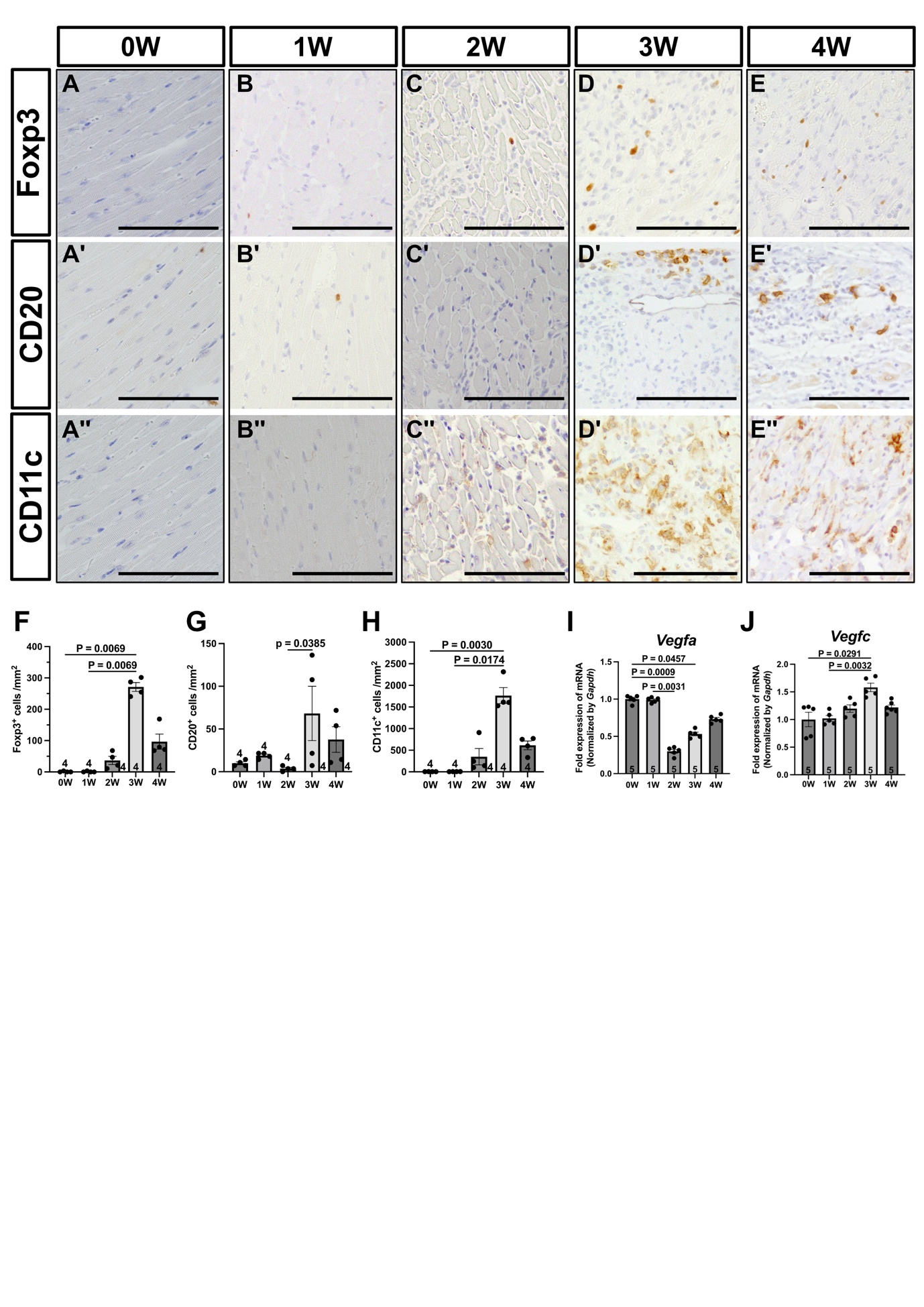
Supplemental Figure 2.

**Supplemental Figure 2. Temporal changes in regulatory T cells, B cells, dendritic cells, and lymphangiogenic gene expression during experimental autoimmune myocarditis.**

(**A–E′′**) Representative immunohistochemistry for Foxp3⁺ regulatory T cells (**A–E**), CD20⁺ B cells (**A′–E′**), and CD11c⁺ cells (**A′′–E′′**) in heart sections collected at 0, 1, 2, 3, and 4 weeks post-immunization. CD11c⁺ cells primarily represent dendritic cells, but may also include activated macrophages. (**F–H**) Quantification of Foxp3⁺, CD20⁺, and CD11c⁺ cells over time.
(**I, J**) Quantitative PCR (qPCR) analysis of *Vegfa* (**I**) and *Vegfc* (**J**) mRNA levels in whole heart tissue at the indicated time points, normalized to *Gapdh*. Each dot represents an individual mouse. Scale bars: 100 μm (**A–E′′**). Statistical analyses were performed using the Kruskal–Wallis test followed by Dunn’s multiple comparison test. Error bars represent the mean ± standard error of the mean (SEM).

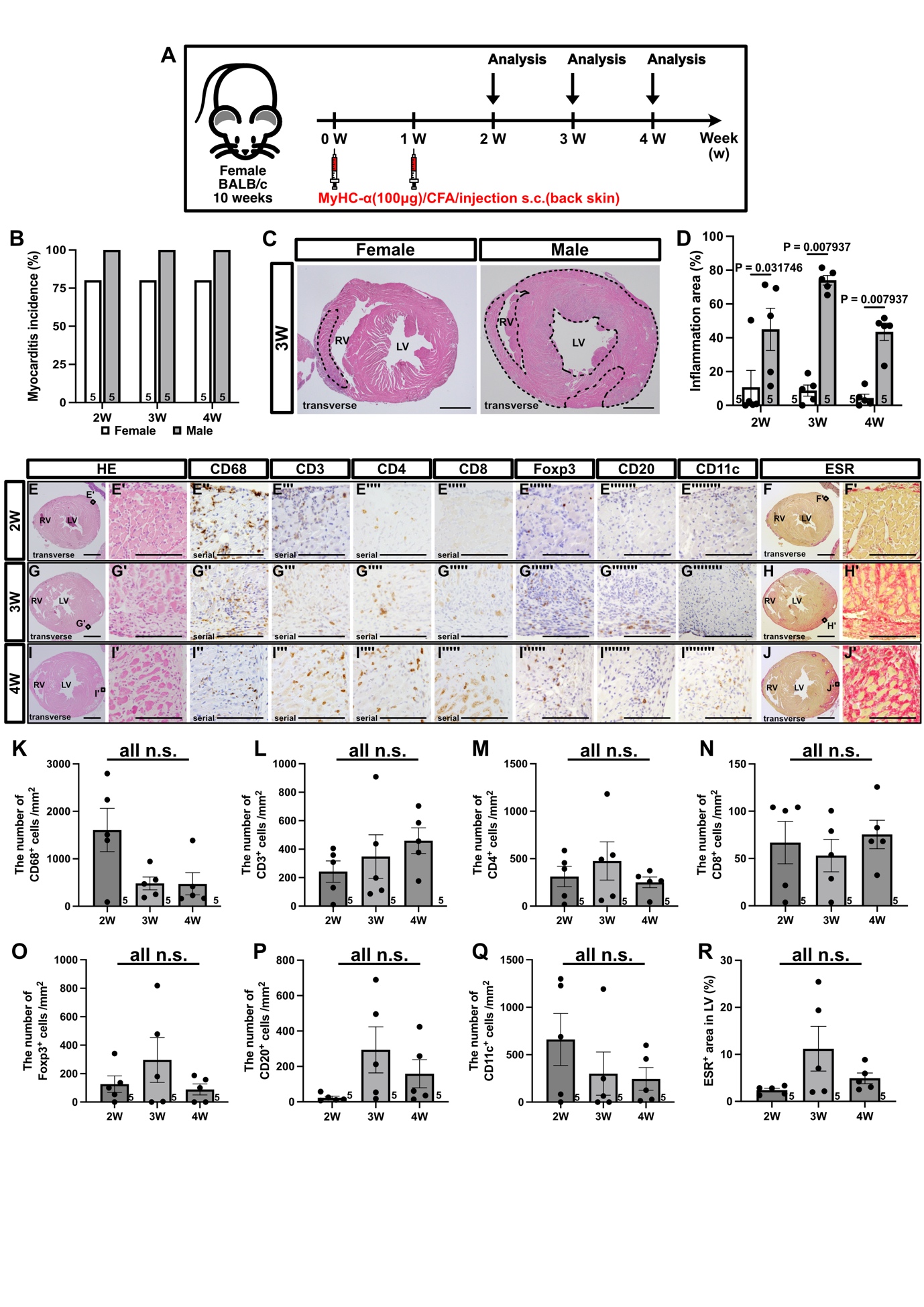
Supplemental Figure 3.

**Supplemental Figure 3. Sex-based differences in myocarditis severity in experimental autoimmune myocarditis.**
(**A**) Experimental timeline for myocarditis induction and analysis in female BALB/c mice. (**B**) Incidence of myocarditis (%) at 2, 3, and 4 weeks post-immunization in female and male mice. (**C**) Representative HE-stained heart sections at 3 weeks, illustrating differences in myocardial inflammation (dotted lines) between sexes. (**D**) Quantification of the inflammation area (% of the total myocardial area, combining the left and right ventricles) in female and male mice. (**E–J′**) Representative histological and immunohistochemical staining in female hearts at 2, 3, and 4 weeks post-immunization, including HE, CD68, CD3, CD4, CD8, Foxp3, CD20, CD11c, and Elastica Sirius Red (ESR). (**K–Q**) Quantification of immune cell infiltration in female mice across the indicated time points. (**R**) Quantification of the fibrotic area (ESR⁺ red-stained region) in the left ventricular (LV) myocardium, showing no significant differences among time points. Each dot represents an individual mouse. Scale bars: 1 mm (**C, E, G, I, F, H, J**) and 100 μm (**E′–E⁗, G′–G⁗, I′–I⁗, F′, H′, J′**). Statistical analyses were performed using the Kruskal–Wallis test followed by Dunn’s multiple comparison test. Error bars represent the mean ± standard error of the mean (SEM).

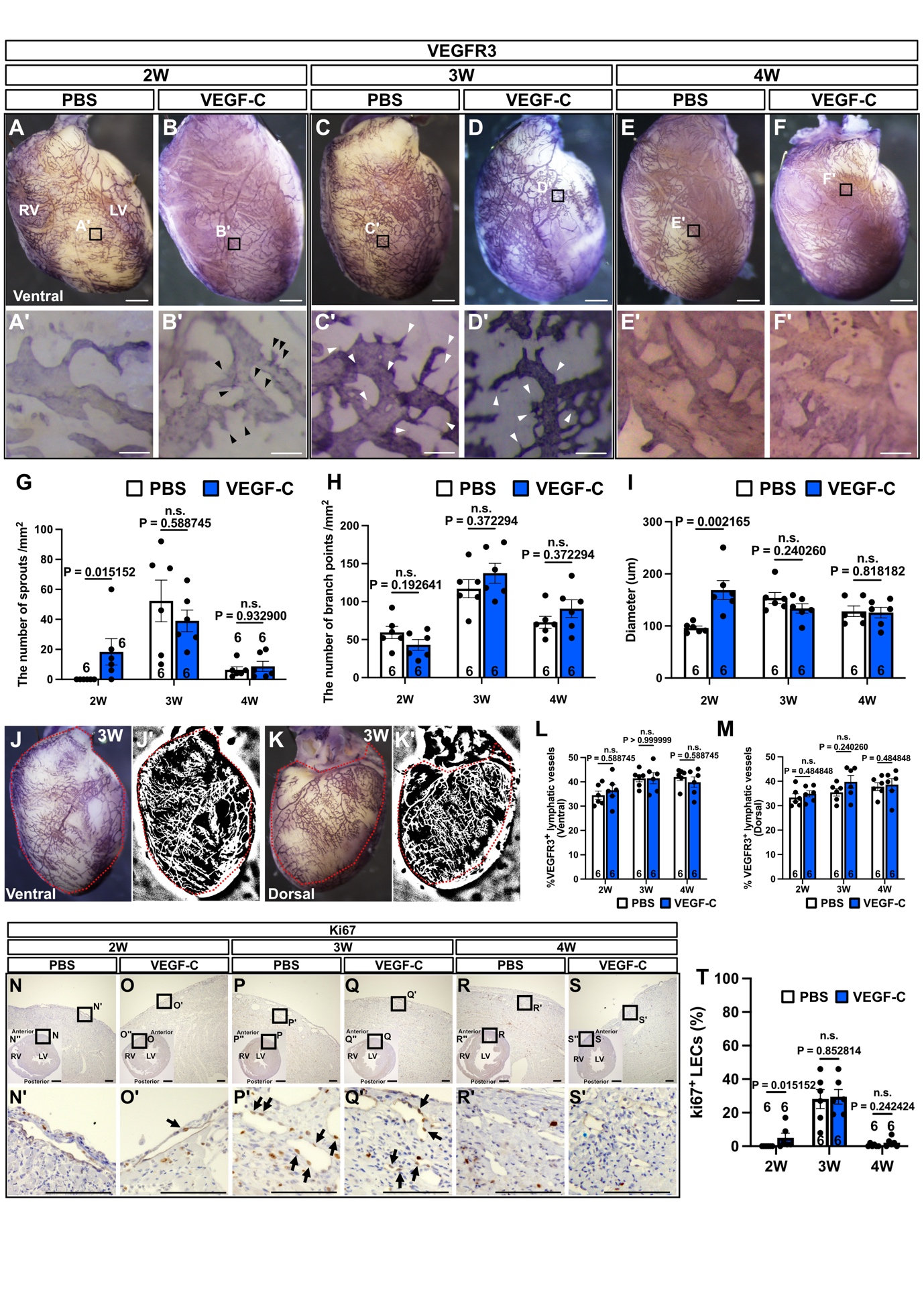
Supplemental Figure 4.

**Supplemental Figure 4. VEGF-C promotes the structural adaptation of cardiac lymphatics during autoimmune myocarditis.**

(**A–F′**) Whole-mount VEGFR3 immunostaining of the ventral heart surface at 2, 3, and 4 weeks post-immunization in PBS- or VEGF-C–treated mice. (**G–I**) Quantification of lymphatic morphological features (sprouts, branch points, and diameters) from whole-mount images. (**J, K**) Representative binarized images of VEGFR3⁺ lymphatic vessels at 3 weeks on the ventral (**J**) and dorsal (**K**) heart surfaces. Red outlines indicate the myocardial area used for quantification. (**L, M**) Quantification of the VEGFR3⁺ lymphatic vessel area (%) on ventral (**L**) and dorsal (**M**) surfaces. (**N–S″**) Paraffin sections prepared after whole-mount staining were subjected to Ki67 immunostaining. VEGFR3⁺ lymphatic structures were developed using a DAB substrate with nickel enhancement, resulting in a purple signal, whereas Ki67 staining appeared brown. (**N–S**) Low-magnification views; (**N′–S′**) enlarged views of boxed regions showing Ki67⁺ lymphatic endothelial cells (black arrows). (**T**) Quantification of proliferating Ki67⁺ LECs, expressed as the percentage of Ki67⁺ nuclei among total LECs. Each dot represents data from an individual mouse. Scale bars: 1 mm (**A–F, N″–S″**) and 100 μm (**A′–F′, N-S, N′–S′**). Statistical analyses were performed using the Mann–Whitney U test. Error bars indicate the mean ± standard error of the mean (SEM).

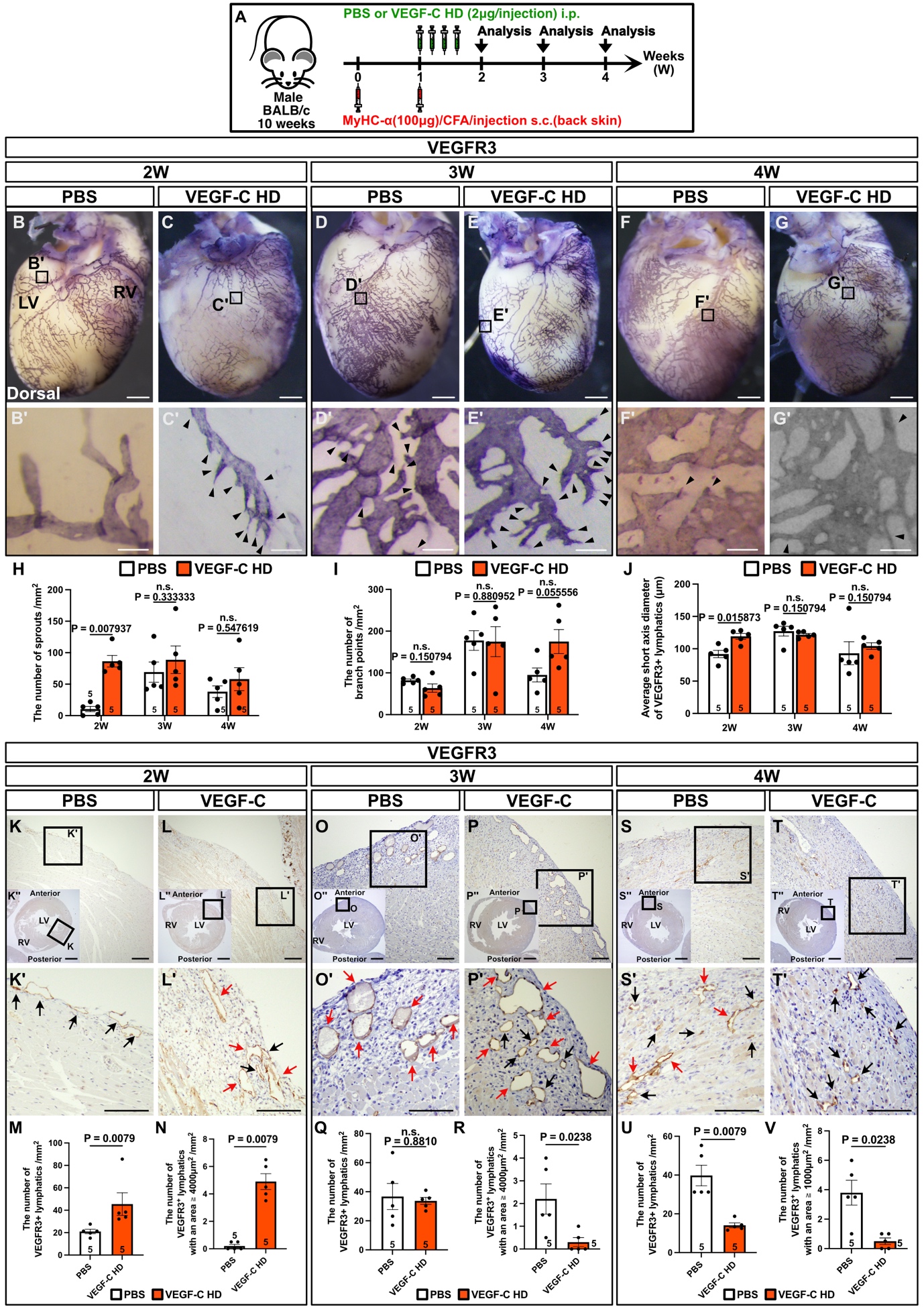
Supplemental Figure 5.

**Supplemental Figure 5. High-dose VEGF-C administration enhances lymphatic sprouting and dilation in experimental autoimmune myocarditis.**

(**A**) Experimental design. Male BALB/c mice (10 weeks old) were immunized with the α-MyHC peptide and CFA on days 0 and 7 to induce myocarditis. Mice were then randomized to receive high-dose VEGF-C C156S (2 μg/injection, intraperitoneally) or PBS daily from Days 7 to 10. Hearts were collected at 2, 3, and 4 weeks post-immunization. (**B–G′**) Representative whole-mount VEGFR3 immunostaining of dorsal heart surfaces at 2, 3, and 4 weeks; black arrowheads indicate lymphatic sprouts. (**H–J**) Quantification of lymphatic sprouting, branching, and vessel diameter from whole-mount images. (**K–L″, O–P″, S–T″**) Representative VEGFR3 immunohistochemistry of paraffin heart sections. (**K–T**) Low-magnification views; (**K′–T′**) higher-magnification views showing VEGFR3⁺ lymphatic vessels (red arrows indicate enlarged lymphatic vessels; black arrows indicate smaller lymphatic vessels). (**M, N, Q, R, U, V**) Quantification of VEGFR3⁺ lymphatic vessels in tissue sections. Each dot represents an individual mouse. Scale bars: 1 mm (**B–G, K″–T″**) and 100 μm (**B′–G′, K–T, K′–T′**). Statistical analyses were performed using the Mann–Whitney U test. Error bars represent the mean ± standard error of the mean (SEM).

Supplemental Figure 6.

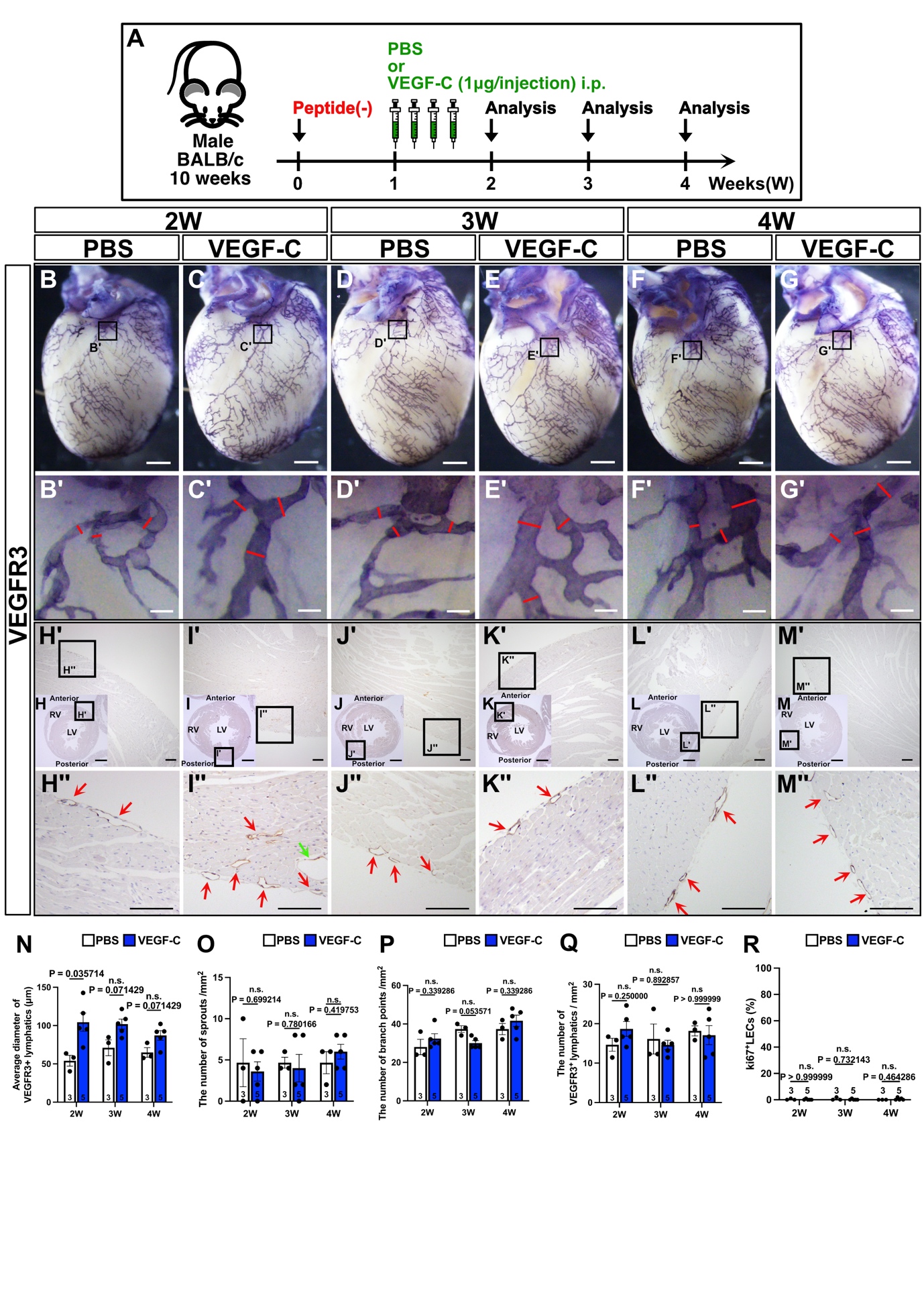

**Supplemental Figure 6. VEGF-C administration does not significantly affect the cardiac lymphatic morphology in non-inflamed hearts.**

(**A**) Experimental design. Male BALB/c mice (10 weeks old) were administered either PBS or VEGF-C C156S (1 μg/injection, intraperitoneally) without myocarditis induction (Peptide(−)). Hearts were collected and analyzed at 2, 3, and 4 weeks post-injection. (**B–G′**) Representative whole-mount VEGFR3 immunostaining of dorsal heart surfaces at 2, 3, and 4 weeks in PBS- or VEGF-C–treated mice. Red lines indicate the short-axis diameters of lymphatic vessels. (**H–M″**) Representative VEGFR3 immunohistochemistry on paraffin heart sections. Red arrows indicate VEGFR3⁺ lymphatic vessels, the green arrow indicates larger VEGFR3^+^ lymphatic vessels. Each dot in the quantification graphs represents an individual mouse. Scale bars: 1 mm (**B–G, H–M**) and 100 μm (**B′–G′, H′–M′, H″–M″**). Statistical analyses were performed using the Mann–Whitney U test. Error bars represent the mean ± standard error of the mean (SEM).

Supplemental Figure 7.

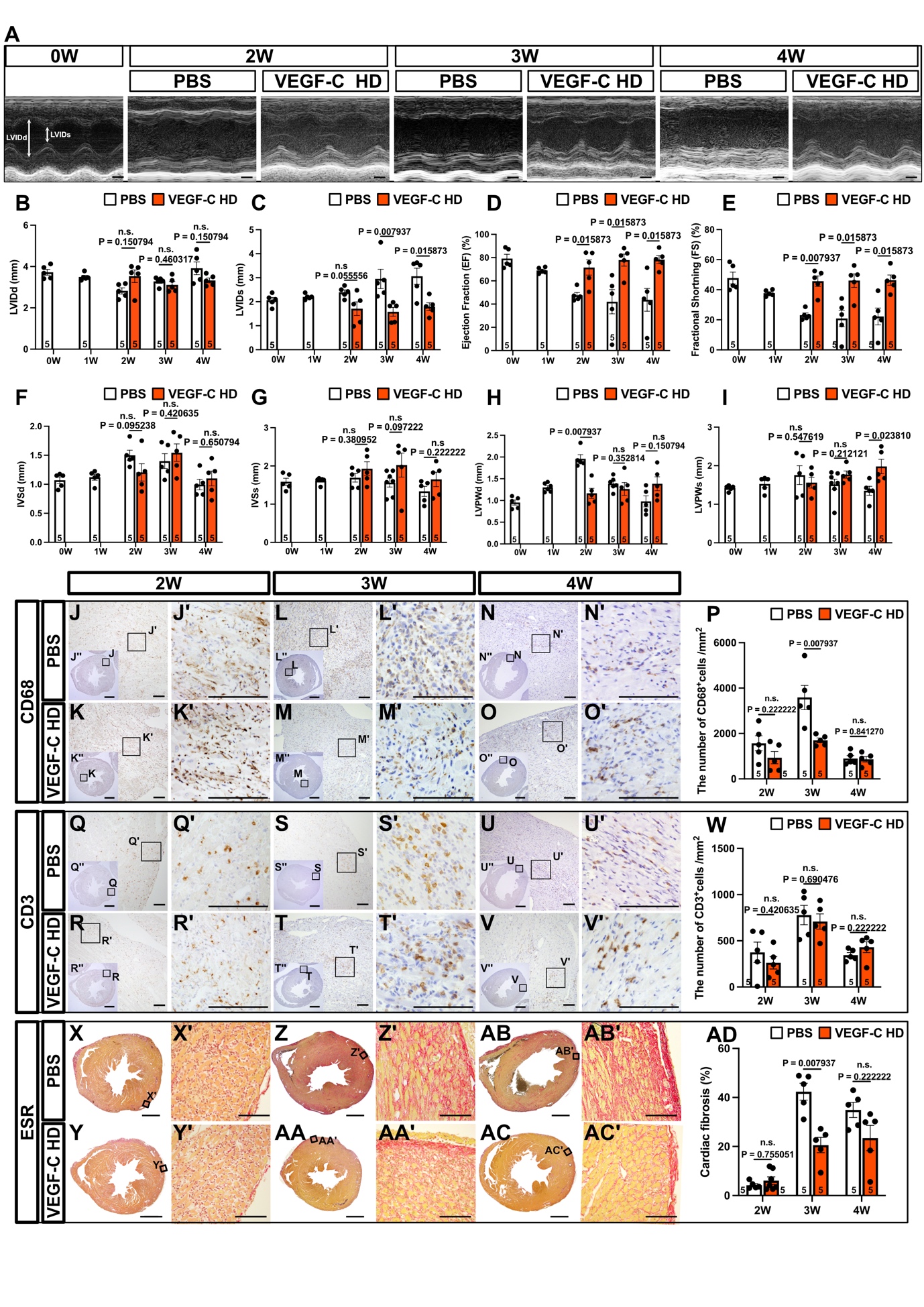

**Supplemental Figure 7. High-dose VEGF-C administration reduces cardiac inflammation and fibrosis in experimental autoimmune myocarditis.**

(**A**) Representative M-mode echocardiographic images of the left ventricle (LV) in PBS- and high-dose VEGF-C (HD)-treated mice at baseline (0W) and at 2, 3, and 4 weeks post-immunization. (**B–I**) Quantification of echocardiographic parameters: (**B**) LVIDd (diastolic LV internal diameter), (**C**) LVIDs (systolic LV internal diameter), (**D**) ejection fraction (EF),
(**E**) fractional shortening (FS), (**F**) interventricular septal thickness at diastole (IVSd), (**G**) at systole (IVSs), (**H**) LV posterior wall thickness at diastole (LVPWd), and (**I**) at systole (LVPWs). (**J–O″**) Representative immunohistochemistry for CD68⁺ macrophages at 2, 3, and 4 weeks post-immunization. (**P**) Quantification of CD68⁺ macrophage density. (**Q–V″**) Representative immunohistochemistry for CD3⁺ T cells at 2, 3, and 4 weeks. (**W**) Quantification of CD3⁺ T cell density. (**X–AC′**) Representative Elastica Picrosirius Red (ESR) staining for cardiac fibrosis (collagen = red) at each time point. (**AD**) Quantification of the fibrotic area (ESR⁺ region as a % of the left ventricular myocardium). Each dot represents data from an individual mouse. Scale bars: 1 mm (**A, J″–O″, Q″–V″, X–AC**) and 100 μm (**J–O, J′–O′, Q–V, Q′–V′, X′–AC′**). Statistical analyses were performed using the Mann–Whitney U test. Error bars represent the mean ± standard error of the mean (SEM).

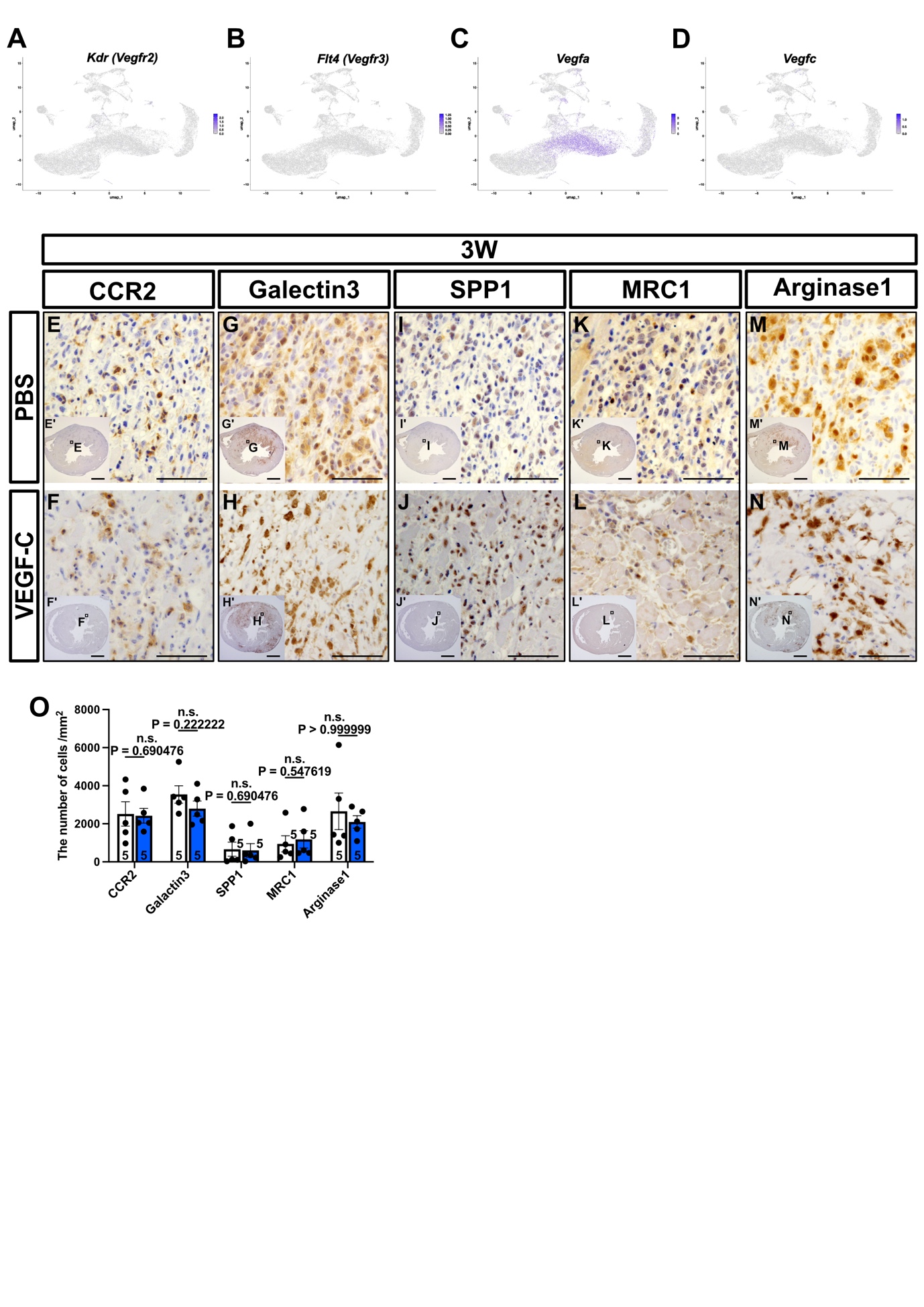
Supplemental Figure 8.

**Supplemental Figure 8. Expression of CCR2, Galectin-3, SPP1, MRC1, and Arginase1 is unchanged by the VEGF-C treatment in autoimmune myocarditis.**

(**A–D**) UMAP feature plots showing the single-cell RNA-seq expression of *Kdr (Vegfr2)*, *Flt4 (Vegfr3)*, *Vegfa*, and *Vegfc* in cardiac CD45⁺ leukocytes isolated from mice with experimental autoimmune myocarditis (EAM). (**E–N′**) Representative immunohistochemistry for CCR2 (**E–F′**), Galectin-3 (**G–H′**), SPP1 **(I–J′**), MRC1 (**K–L′**), and Arginase1 (**M–N′**) in hearts from PBS- and VEGF-C–treated mice at 3 weeks post-immunization. (**O**) Quantification of the number of positive cells for each marker. No significant differences were observed between PBS- and VEGF-C–treated groups. Each dot represents data from an individual mouse. Scale bars: 1 mm (**E′–N′**) and 100 μm (**E–N**). Statistical analyses were performed using the Mann–Whitney U test. Error bars represent the mean ± standard error of the mean (SEM).
